# Large-scale computational discovery and analysis of virus-derived microbial nanocompartments

**DOI:** 10.1101/2021.03.18.436031

**Authors:** Michael P. Andreas, Tobias W. Giessen

**Affiliations:** Department of Biomedical Engineering, University of Michigan Medical School, Ann Arbor, MI, USA; Department of Biological Chemistry, University of Michigan Medical School, Ann Arbor, MI, USA

## Abstract

Protein compartments represent an important strategy for subcellular spatial control and compartmentalization. Encapsulins are a class of microbial protein compartments defined by the viral HK97-fold of their capsid protein, self-assembly into icosahedral shells, and dedicated cargo loading mechanism for sequestering specific enzymes. Encapsulins are often misannotated and traditional sequence-based searches yield many false positive hits in the form of phage capsids. This has hampered progress in understanding the distribution and functional diversity of encapsulins. Here, we develop an integrated search strategy to carry out a large-scale computational analysis of prokaryotic genomes with the goal of discovering an exhaustive and curated set of all HK97-fold encapsulin-like systems. We report the discovery and analysis of over 6,000 encapsulin-like systems in 31 bacterial and 4 archaeal phyla, including two novel encapsulin families as well as many new operon types that fall within the two already known families. We formulate hypotheses about the biological functions and biomedical relevance of newly identified operons which range from natural product biosynthesis and stress resistance to carbon metabolism and anaerobic hydrogen production. We conduct an evolutionary analysis of encapsulins and related HK97-type virus families and show that they share a common ancestor. We conclude that encapsulins likely evolved from HK97-type bacteriophages. Our study sheds new light on the evolutionary interplay of viruses and cellular organisms, the recruitment of protein folds for novel functions, and the functional diversity of microbial protein organelles.

## Introduction

Spatial compartmentalization is a ubiquitous feature of biological systems.^1^ In fact, biological entities like cells and viruses only exist because of the presence of a barrier that separates their interior from the environment. This concept of creating distinct spaces separate from their surroundings extends further to intracellular organization with many layers of sub-compartmentalization found within most cells.^2,3^ Intracellular compartments with a proteomically defined interior and a discrete boundary that fulfill distinct biochemical or physiological functions are generally referred to as organelles.^4^ This includes both lipid-bound organelles, phase-separated structures, and protein-based compartments. Distinguishing features between eukaryotic lipid-based and prokaryotic protein-based organelles include their size range – micro vs. nano scale – and the fact that protein organelle structure is genetically encoded and thus generally more defined. Still, compartmentalization, however it is achieved, can ultimately serve four distinct functions, namely, the creation of distinct reaction spaces and environments, storage, transport, and regulation.^4^ Often, compartmentalization can serve multiple of these functions at the same time. More specifically, the functions of intracellular compartments include sequestering toxic reactions and metabolites, creating distinct biochemical environments to stimulate enzyme or pathway activity, and dynamically storing nutrients for later use, among many others.^4^

One of the most widespread and diverse classes of protein-based compartments are encapsulin nanocompartments, or simply encapsulins.^5-7^ So far, two families of encapsulins have been reported in a variety of bacterial and archaeal phyla.^8-10^ They are proposed to be involved in oxidative stress resistance,^9,11-13^ iron mineralization and storage,^14,15^ anaerobic ammonium oxidation,^16^ and sulfur metabolism.^8^ All known encapsulins self-assemble from a single capsid protein into compartments between 24 and 42 nm in diameter with either T=1, T=3 or T=4 icosahedral symmetry.^10,12,15^ Their defining feature is the ability to selectively encapsulate cargo proteins which include ferritin-like proteins, hemerythrins, peroxidases and desulfurases.^8,9^ In classical encapsulins (Family 1), encapsulation is mediated by short C-terminal peptide sequences referred to as targeting peptides (TPs) or cargo-loading peptides (CLPs)^10,15,17^ while for Family 2 systems, larger N-terminal protein domains are proposed to mediate encapsulation.^8^ For most encapsulin systems, little is known about the specific reasons or functional consequences of enzyme encapsulation. Suggestions include the sequestration of toxic or reactive intermediates as well as enhancing enzyme activity and the prevention of unwanted side reactions. One of the most intriguing features of encapsulins is that in contrast to all other known protein-based compartments or organelles, their capsid monomer shares the HK97 phage-like fold.^10,12,15^ This has led to the suggestion that encapsulins are derived from or in some way connected to the world of phages and viruses.^5,9^

Here, we carry out a large-scale in-depth computational analysis of prokaryotic genomes with the goal of discovering and classifying an exhaustive set of all HK97-type protein organelle systems. We develop a Hidden Markov Model (HMM)-, Pfam family-, and genome neighborhood analysis (GNA)-based search strategy and substantially expand the number of identified encapsulin-like operons. We report the discovery and analysis of two novel encapsulin families (Family 3 and Family 4) as well as many new operon types that fall within Family 1 and Family 2. We formulate data-driven hypotheses about the potential biological functions of newly identified operons which will guide future experimental studies of encapsulin-like systems. Further, we conduct a detailed evolutionary analysis of encapsulin-like systems and related HK97-type virus families and show that encapsulins and HK97-type viruses share a common ancestor and that encapsulins likely evolved from HK97-type phages. Our study sheds new light on the evolutionary interplay of viruses and cellular organisms, the recruitment of protein folds for novel functions, and the functional diversity of microbial protein organelles.

## Results and Discussion

### Distribution, diversity, and classification of encapsulin systems found in prokaryotes

All bacterial and archaeal proteomes available in the UniProtKB^18^ database (Family 1, 2, and 4: March 2020; Family 3: February 2021) were analyzed for the presence of encapsulin-like proteins using an HMM-based search strategy. It was discovered that all Pfam families associated with initial search hits belong to a single Pfam clan (CL0373)^19^ encompassing the majority of HK97-fold proteins catalogued in the Pfam database. Thus, we supplemented our initial hit dataset with all sequences associated with CL0373. This was followed by GNA-based curation^20^ of the expanded dataset to remove all false positives, primarily phage genomes, resulting in a curated list of 6,133 encapsulin-like proteins (**Fig. 1** and **Supplementary Data 1**). Encapsulin-like systems can be found in 31 bacterial and 4 archaeal phyla. Based on the sequence similarity and Pfam family membership of identified capsid proteins, and the genome-neighborhood composition of associated operons, encapsulin-like systems could be classified into 4 distinct families (**Fig. 2**). Family 1 and 2 represent previously identified encapsulin operon types containing capsid proteins falsely annotated as *bacteriocin* (PF04454: Linocin_M18) and *transcriptional regulator/membrane protein* (no Pfam), respectively. Family 1 will be referred to as Classical Encapsulins given the fact that they were the first discovered and are the best characterized. Family 3 and 4 represent newly discovered systems. Family 3 encapsulins are falsely annotated as *phage major capsid protein* (PF05065: Phage_capsid) and are found embedded within large biosynthetic gene clusters (BGCs) encoding different peptide-based natural products. Therefore, Family 3 was dubbed Natural Product Encapsulins. Family 4 is characterized by a highly truncated encapsulin-like capsid protein which is generally annotated as an *uncharacterized protein* (PF08967: DUF1884) and arranged in conserved two-component operons with different enzymes. Family 4 proteins represent the A-domain of the canonical HK97-fold with all other domains usually associated with this fold missing. Thus, Family 4 will be referred to as A-domain Encapsulins.

**Fig. 1.**
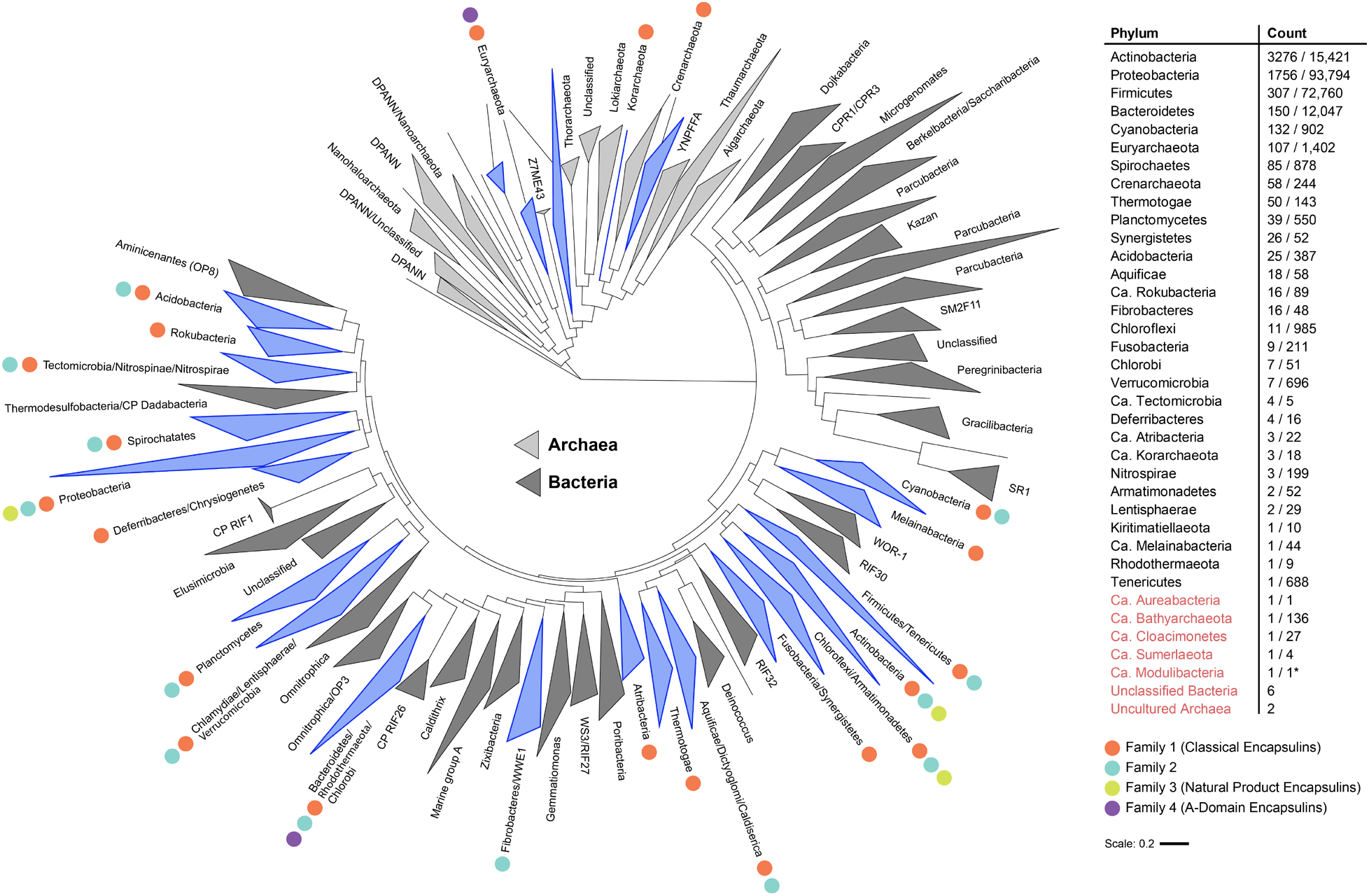
Distribution of encapsulin-like systems in prokaryotes. Left: Phylogenetic tree based on 108 of the major archaeal and bacterial phyla.^21^ Phyla containing encapsulin-like systems are highlighted in blue. Differently colored dots indicate the presence of the respective encapsulin family within the phylum. Right: List of phyla discovered to encode encapsulin-like systems. The *Count* column shows the number of identified systems and the total number of proteomes available in UniProt (# systems identified / # UniProt proteomes). Ca. refers to candidate phyla. Phylum names colored red show new phyla or uncultured/unclassified organisms not shown in the phylogenetic tree. *Ca. Modulibacteria is not an annotated phylum in UniProt but has been proposed as a candidate phylum.^22^

**Fig. 2.**
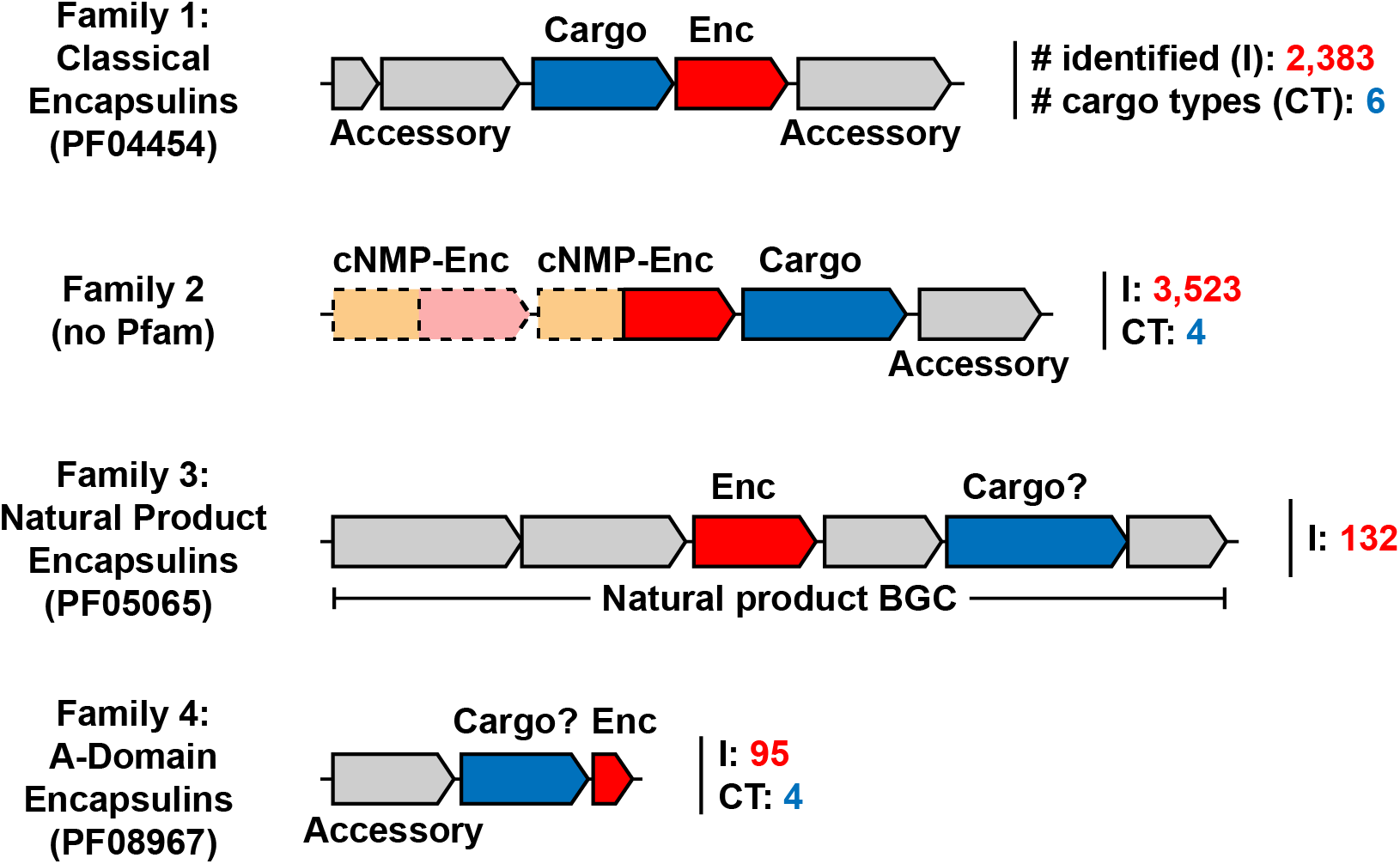
Novel classification scheme for encapsulin-like operons. Shown are the 4 newly defined families of encapsulins with the respective Pfam annotations if available. Encapsulin-like capsid components are shown in red. Confirmed and proposed cargo proteins are shown in blue. Non-cargo accessory components are shown in grey. The number of identified systems of a given family is shown after the operon in red (I, # identified) and the number of distinct cargo types is shown in cyan (CT, # cargo types). Dotted lines indicate optional presence of operon components. cNMP: cyclic nucleotide-binding domain (orange), Enc: encapsulin-like capsid component. BGC: biosynthetic gene cluster.

Classical Encapsulins (Family 1) represent the most widespread family of encapsulin-like systems. They can be found in 31 out of 35 prokaryotic phyla found to encode encapsulin-like operons (**Fig. 1**). 2,383 Classical Encapsulin operons were discovered with the phyla Proteobacteria, Actinobacteria and Firmicutes containing the majority of identified systems. However, it should be noted that these phyla also contain the largest number of sequenced genomes and available proteomes. Family 1 contains at least 6 operon types defined by the presence of 6 distinct and conserved cargo proteins. Many of these operon types can be found in distantly related phyla consistent with frequent horizontal gene transfer events. The general operon organization of Family 1 systems consists of the encapsulin capsid protein and a single primary cargo protein usually encoded directly upstream of the shell component (**Fig. 2**).^9^ Depending on the operon type, other conserved accessory components can be present.^9,15^ These components are not cargo proteins but are proposed to be directly involved in the biochemical function or regulation of a given system.

Family 2 encapsulins are the most numerous encapsulin-like systems and can be found in 14 bacterial phyla (**Fig. 1**). 3,523 Family 2 operons were identified. The majority of systems can be found in the phyla Actinobacteria and Proteobacteria followed by Bacteroidetes and Cyanobacteria. Family 2 contains at least 4 different operon types based on cargo protein identity. Again, the widespread occurrence of these operon types in distant phyla supports the hypothesis of their frequent horizontal transfer. Family 2 operon organization is more complex compared to Family 1 due to the variable presence of a cNMP-binding domain (PF00027) fused to the encapsulin capsid component as well as the variable occurrence of two distinct capsid components within a single Family 2 operon. Further non-cargo accessory components may be present, likely related to the biological function of a given operon (**Fig. 2**).^8^

Natural Product Encapsulins (Family 3) can be found almost exclusively in the phyla Actinobacteria and Proteobacteria, primarily in *Streptomyces* and *Myxococcus* species as well as some other closely related genera (**Fig. 1**). *Streptomyces* and *Myxococcus* species are widely known as being among the most prolific producers of bioactive natural products.^23,24^ So far, 132 Family 3 systems have been identified and can be classified into 6 distinct operon types based on the organization of the BGC surrounding the encapsulin capsid component (**Fig. 2**).

A-domain Encapsulins (Family 4) are the most distinct family so far discovered and are restricted to the archaeal phylum Euryarchaeota and the bacterial phylum Bacteroidetes (**Fig. 1**). 95 Family 4 operons have been identified with more than 90 percent found in Archaea. Family 4 encapsulin-like proteins are truncated and thus only one third the length of a standard HK97-fold protein.^25,26^ All archaeal Family 4 operons consist of a single- or multi-subunit enzyme and the A-domain Encapsulin protein located downstream of the enzymatic component (**Fig. 2**). Some systems seem to possess further accessory components as part of the operon as judged by overlapping genes and transcription direction. So far, 4 distinct archaeal operon types have been discovered. The identified bacterial A-domain Encapsulins are not arranged in an obvious operon-like structure which makes their classification and function prediction more difficult.

### Family 1 – Classical Encapsulins

Our dataset of 2,383 Family 1 systems greatly expands the set of the previously described 932 Classical Encapsulins (**Fig. 3**).^9,16^ 1,505 dye-decolorizing peroxidase (DyP) systems were identified, making them the most abundant cargo class in Family 1. DyP systems are most abundant in *Actinobacteria* and *Proteobacteria*, with 962 and 519 systems found in each phylum, respectively. DyP peroxidases bind heme and are named for their ability to oxidize a broad range of anthraquinone dyes.^27^ DyPs have also been shown to break down lignin and other typical peroxidase substrates.^28,29^

**Fig. 3.**
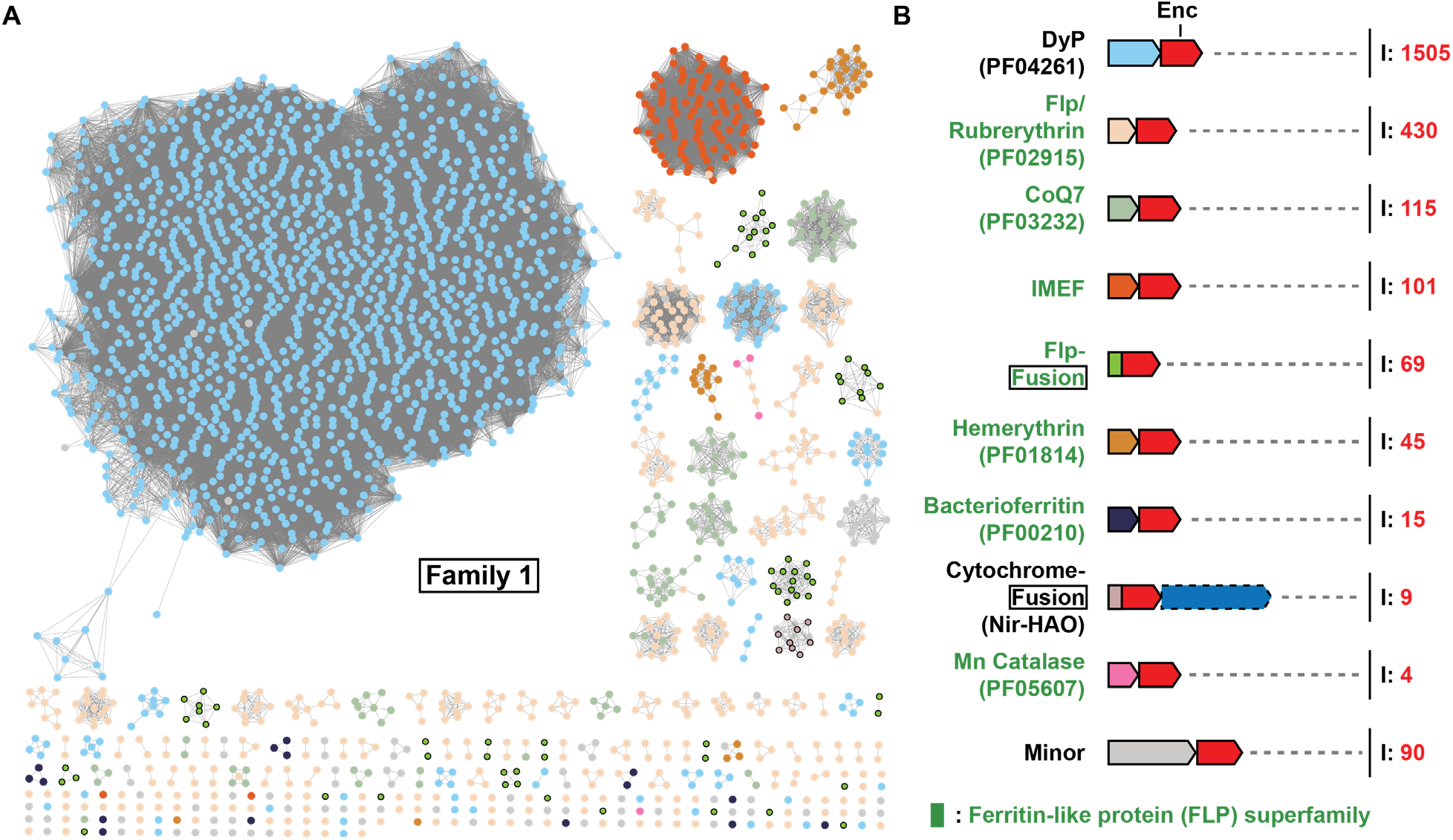
Overview and analysis of Family 1 encapsulin systems. A) SSN analysis of 2383 Family 1 encapsulins clustered at 49% sequence identity. Nodes are colored based on the associated primary cargo type shown in B). B) Diversity of Family 1 operon types. Only conserved primary cargo proteins are shown. Operons not containing any of the main cargo types are designated as Minor and are shown in more detail in **Fig. S1**. I: number of identified operons.

Encapsulated DyP from *Brevibacterium linens* has been shown to form a trimer of dimers with D3 symmetry and bind close to the three-fold symmetry axis of the encapsulin shell via C-terminal targeting peptides.^10^ Many DyP Family 1 operons in *Mycobacteria* contain accessory genes encoding short chain oxidoreductases and cupins in addition to the core DyP cargo. Their function within the context of DyP encapsulin operons is currently unknown (**Fig. S1A**). Accessory genes encoding putative membrane proteins containing DUF1345 domains are commonly found in DyP-containing operons in *Streptomyces* and might play a role in transport related to DyP function (**Fig. S1A**). Further, 67 DyP operons were identified in *Streptomyces* that contain accessory genes encoding for a DUF5709 domain protein and genes annotated as 6-phosphogluconate dehydrogenases (6-GPD) and diaminopimelate decarboxylases (DAPDC) (**Fig. S1A**). Both 6-GPD and DAPDC possess decarboxylase activity and play key roles in the pentose phosphate pathway and amino acid biosynthesis, however, their role in the context of DyP encapsulin systems is currently unknown. The general biological function of DyP encapsulin systems is still speculative, however, a recent study showed that a DyP Family 1 system in *Mycobacterium tuberculosis* plays a direct role in oxidative stress resistance during infection.^11^

Ferritin-like proteins (FLPs) comprise the second largest set of cargo proteins associated with Family 1 encapsulins. FLPs represent a large functionally diverse superfamily of proteins that all share a four-helix bundle fold. Clustering encapsulin-associated FLPs at 30% sequence identity results in 7 distinct families that largely correspond to the following Pfam families: Flp (lower case to distinguish the Pfam family from the superfamily), rubrerythrin, CoQ7, Mn catalase, IMEF, hemerythrin, and bacterioferritin (**Fig. S2**).

Identified Flp, rubrerythrin, CoQ7, and Mn catalase cargo proteins are likely functionally identical – acting as ferroxidases – and should likely be part of the same Pfam family. From now on, we will refer to all four simply as Flp cargos. They are found in 23 bacterial and 2 archaeal phyla. They are widespread in bacteria but predominantly found in *Firmicutes* and *Proteobacteria*. Crystal structures of encapsulin-associated Flps from *Haliangium ochraceum* and *Rhodospirillum rubrum* suggest that these systems form decameric assemblies with D5 symmetry (**Fig. S2**).^14,30^ Unlike ferritin cages with higher symmetries, Flp cargo proteins cannot store precipitated iron in a soluble form by themselves and rely on the encapsulin shell to achieve iron precipitate sequestration. Similar to the ubiquitous ferritin iron storage cages, Flp encapsulin systems might play a dual role in oxidative stress resistance and iron homeostasis.

The second largest cargo class within the FLP superfamily are the iron-mineralizing encapsulin-associated Firmicute (IMEF) cargos. They form dimers in solution and when encapsulated and are most commonly found in *Firmicutes*. Encapsulins containing these systems form large T=4 capsids approximately 42 nm in diameter.^9,15^ The large size of these assemblies allows them to form iron-rich cores up to 30 nm in diameter, making them the largest protein-based iron storage system known to date. Many IMEF-containing operons encode 2Fe-2S ferredoxins (Fdxs) homologous to bacterioferritin-associated ferredoxins (Bfds) (**Fig. S1**). Bfd proteins assist in the mobilization of iron from iron-filled ferritin cages.^31^ Many of the identified Fdxs contain a strongly conserved targeting peptide-like TVGSL motif at their N-terminus and have been shown to co-purify with IMEF encapsulins when heterologously expressed.^9^ Fdxs might be involved in releasing stored iron from IMEF encapsulins by transferring electrons to the interior of the capsid, thus reducing and solubilizing stored iron. Most organisms encoding IMEF systems do not encode any classical ferritins making it likely that IMEF encapsulins act as their primary iron storage compartments.

Within the FLP superfamily, 45 hemerythrin cargos were identified, with 42 found in *Actinobacteria* and 3 in *Proteobacteria*. No hemerythrin-containing encapsulin has been structurally characterized, but hemerythrins have been shown to form dimers in solution.^9^ Hemerythrin cargos have further been shown to offer oxidative and nitrosative stress protection when encapsulated.^9^ All hemerythrins contain binuclear iron centers which have been shown to bind to nitric oxide, oxygen, and other reactive or volatile small molecules.^32,33^ Family 1 hemerythrin systems are thus likely involved in the sequestration and detoxification of harmful compounds.

Another FLP cargo type identified in a small number of *Firmicutes, Aquificae, Chlorflexi*, and *Cyanobacteria* are bacterioferritins (Bfrs). These putative cargos are composed of two four-helix bundles and are thus structurally distinct from the other identified FLP superfamily cargos (**Fig. S2**). Bfrs generally assemble into 24 subunit 12 nm cages able to store iron – similar to eukaryotic ferritins.^34^ A bacterioferritin (BfrB) encoded outside a Family 1 operon has been proposed to be a potential cargo protein in *M. tuberculosis*, however, no Family 1 operon encoding a Bfr cargo protein has been reported before.^13^ The presence of conserved C-terminal targeting peptides in the identified Bfrs strongly suggests that they are encapsulin cargos. The biological function and underlying logic of a putative shell-within-a-shell arrangement in the context of iron storage compartments is currently unknown.

The number of Flp-fusion encapsulins was also expanded. In these systems, an Flp domain is N-terminally fused to the encapsulin capsid protein. This leads to the internalization of Flp domains upon capsid self-assembly. All Flp-fusion systems are present in Archaea, mostly in the phylum *Crenarchaeota*. Structural studies of *Pyrococcus furiosus* and *Sulfolobus solfataricus* Flp-fusion encapsulins have shown that these systems assemble into T=3 capsids and contain internalized Flp assemblies.^35,36^ While the excised *P. furiosus* Flp domain has been shown to form a decamer with D5 symmetry – similar to other characterized Flp cargos – the structural arrangement of fused and encapsulated Flps remains unknown.^14^ Flp-fusion encapsulins are often located in operons containing other ferritin-like proteins or rubrerythrins, hinting at a function related to iron homeostasis and stress resistance (**Fig. S1**).

Another type of Family 1 encapsulin fusion system was identified in *Planctomycetes*. 9 encapsulin-encoding genes with an N-terminal diheme cytochrome C fusion domain were identified. All cytochrome-fusion systems are found in anammox bacteria and are associated with a nitrite reductase-hydroxylamine oxidoreductase (NIR-HAO)-encoding gene. These systems have been shown to form T=3 icosahedral compartments.^9^ Their biological function is currently unknown, however, a role in detoxifying harmful intermediates of the anammox process like nitric oxide, hydroxylamine, and hydrazine, has been proposed, as well as a role in iron storage inside the anammoxosome – the membrane-bound compartment sequestering the anammox process in anammox bacteria.^9,16^

90 Family 1 systems were identified which could not be assigned to any of the so far discussed operon types. They are present in a broad range of phyla and are found within diverse genome neighborhoods (**Fig. S1**). Of note are a number of systems found in S*treptomyces* and *Mesorhizobium* species with conserved N-terminal truncations of the encapsulin capsid gene which might indicate a divergent mode of capsid assembly (**Fig. S1B**).

### Family 2

Family 2 encapsulins are the most abundant class of encapsulins identified in this study and can be broadly grouped into two structurally distinct variants: Family 2A and Family 2B. A classification of Family 2 systems was previously proposed based on phylogeny and cargo protein type.^8^ However, with our more expanded dataset, it became clear that a classification based on the most distinctive feature of this class, namely, the absence (2A) or presence (2B) of an internal cNMP-binding domain, would be more appropriate. All Family 2 encapsulins display the HK97-like fold but do not contain an elongated N-terminal helix seen in Family 1 encapsulins (**Fig. S3**). Instead, they possess an extended N-arm with a short N-terminal α-helix (N-helix), more characteristic of the canonical HK97 fold found in bacteriophages.^8^ A single Family 2A structure has been solved (*Synechococcus elongatus*) (PDB: 6×8M and 6×8T)^8^ which showed that the N-terminus, including the N-helix, extends towards the outside of the capsid, in contrast to Family 1 encapsulins where the N-terminus is sequestered inside the protein shell. Family 2 encapsulins generally contain an extended N-terminal sequence (N-extension) preceding the N-helix (**Fig. S3**). Family 2A encapsulins display short N-extensions around 11 amino acids long while Family 2B encapsulins tend to have longer N-extensions around 18 amino acids in length. All N-extensions are predicted to be disordered. In the extreme case of the *Streptomyces parvulus* Family 2 system, the N-extension is 88 amino acids long. The putative cNMP-binding domains in Family 2B encapsulins are also highly variable, sharing only 19% pairwise identity between all identified domains. The cNMP-binding domain is connected to the C-terminal fragment of the E-loop via a poorly conserved ca. 60 amino acid linker that is predicted to be disordered (**Fig. S3**). The presence of the cNMP-binding domain suggests that Family 2B encapsulins may regulate encapsulated components in a cNMP-dependent manner, providing these systems with a novel mode of enzyme regulation via sequestration inside a protein shell, not seen in any other encapsulin family.

The most commonly enriched genes associated with Family 2 encapsulins encode for cysteine desulfurases (CD), terpene cyclases (TC), polyprenyl transferases (PT), and xylulose kinases (XK) (**Fig. 4A** and **4B**). With the exception of xylulose kinases, all of these genes encode proteins with large unannotated regions at their termini, generally predicted to be disordered, which may be involved in mediating cargo encapsulation (**Fig. 4C** and **Fig. S4**). Family 2B operons often encode two distinct cNMP-domain-containing encapsulin capsid proteins (**Fig. 4B**). This opens up the intriguing possibility of encapsulins forming two-component shells. Family 2B capsids encoded within the same operon roughly share 60% sequence identity with the main differences being primarily found in the E-loops and putative cNMP-binding domains. This relatively low sequence identity, the localized sequence differences, and the conservation of double shell systems across many phyla and cargo types likely means that encoding two capsid proteins in a single operon is a feature of these systems and not the result of a recent gene duplication event. The presence of two distinct regulatory cNMP-binding domains within the same capsid may allow the fine-tuning of the activity of encapsulated cargo. However, we can currently not exclude that these operons encode two separately assembling encapsulins instead of a single mixed shell.

**Fig. 4.**
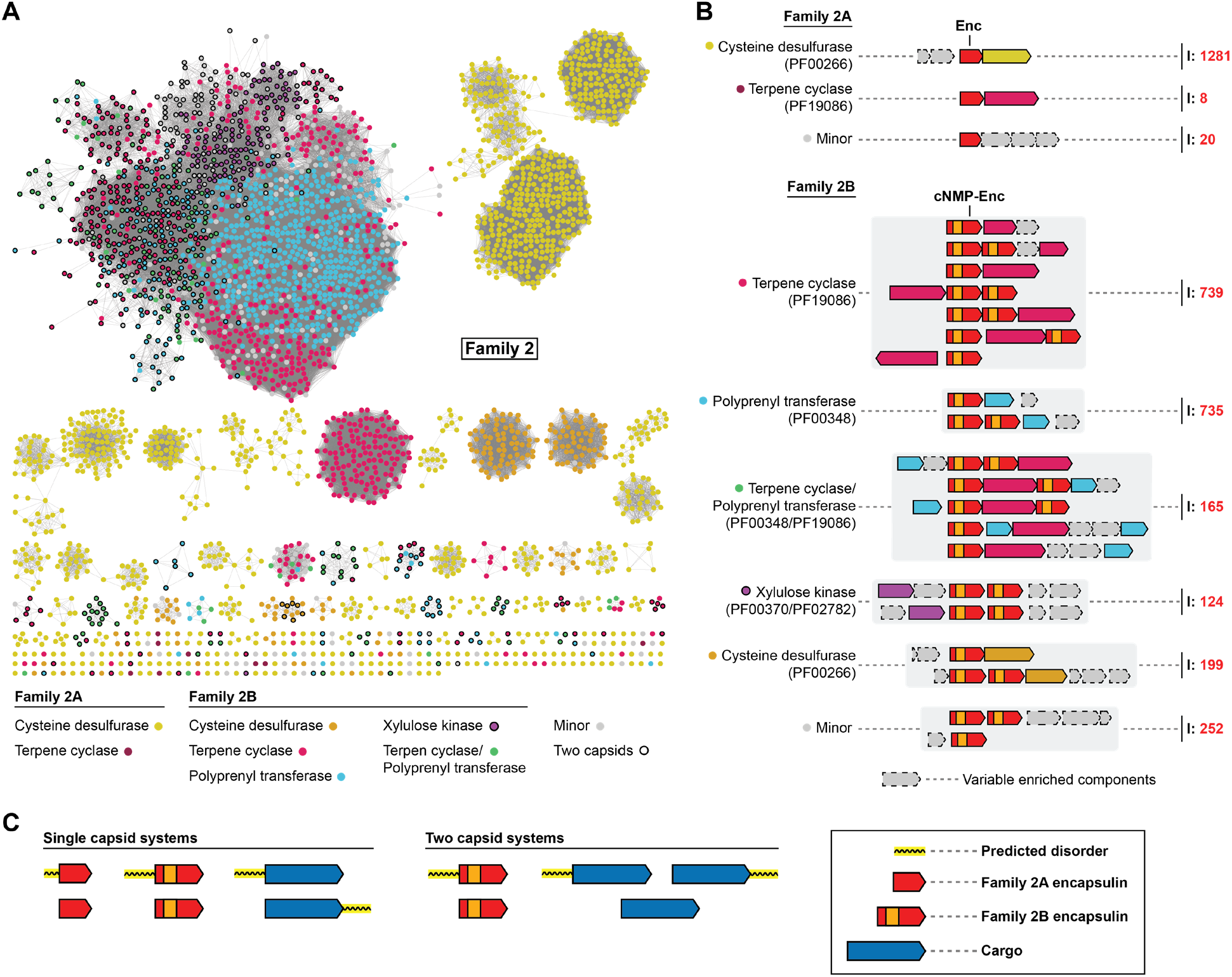
Overview and analysis of Family 2 encapsulin systems. A) SSN analysis of 3523 Family 2 encapsulins clustered at 70% sequence identity. Nodes are colored based on the putative associated cargo type. Family 2A: no cNMP domain, Family 2B: cNMP domain present. B) Selection of operon types encoding Family 2 encapsulins. Operons are grouped by their conserved putative cargo protein type. C) Combinations of commonly observed extended disordered regions at the termini of Family 2 encapsulins and associated cargo proteins. I: number of identified operons.

The partially characterized *S. elongatus* Family 2A encapsulin has been shown to encapsulate a CD cargo protein and to be upregulated during sulfur starvation.^8^ We have identified 1,281 Family 2A and 199 Family 2B encapsulin-encoding operons containing CDs as the putative cargo (**Fig. 4**). Family 2A CD systems are present in 12 bacterial phyla and are most abundant in Proteobacteria (813), Actinobacteria (193), and Bacteroidetes (111). Family 2B CD systems can be found in 9 phyla with a similar distribution as Family 2A systems. The N-termini of CDs are largely predicted to be disordered and are not annotated while the C-terminal region contains a conserved SufS-like cysteine desulfurase domain (PF00266) (**Fig. S4**) that usually converts cysteine to alanine whilst using the liberated sulfur atom to form a protein-bound persulfide intermediate which is then transferred to sulfur acceptor proteins.^37^ While no specific targeting peptide has been identified in CD systems, the unannotated N-terminal domain has been shown to be responsible for mediating encapsulation.^8^ Serine *O*-acetyltransferases and rhodaneses are the most highly enriched accessory components found in these operons (**Fig. S5**). Serine *O*-acetyltransferases catalyze the formation of *O*-acetyl-serine, which is then converted to cysteine via cysteine synthase. Rhodaneses typically act as sulfur atom acceptors, distributing sulfur to various metabolic pathways and processes including cofactor biosynthesis and iron-sulfur cluster formation.^38^ Sequestering a CD inside a protein shell might ensure that only a specific co-regulated rhodanese able to interact with the encapsulin capsid exterior can act as the sulfur acceptor thus making sure that sulfur is channeled to a specific subset of metabolic targets. The presence of these operon components suggests that Family 2A CD systems play a role in sulfur utilization and redox homeostasis.

TC-and PT-encoding genes are highly enriched in many Family 2B operons suggesting a role in terpenoid biosynthesis (**Fig. 4**). We have identified 904 Family 2B operons encoding TCs as their putative cargo. They are commonly found in Actinobacteria (724), Proteobacteria (114), and Cyanobacteria (64). PT systems were found in 900 operons – almost exclusively in Actinobacteria (888). 165 systems were found to encode both TCs and PTs in the same gene cluster. The operon structure of these systems is highly diverse. Many TC systems encode *C*-methyltransferases, usually associated with 2-methylisoborneol-synthase (2-MIBS)-like TCs. Isopentenyl pyrophosphate isomerases and alcohol dehydrogenases are also enriched in TC operons and likely add to the diversity of terpenoid products produced by these systems (**Fig. S5**). PT systems often encode genes involved in terpenoid precursor biosynthesis. Other genes enriched in PT operons encode terpenoid tailoring enzymes like epimerases, dehydrogenases, acetyltransferases, and deaminases indicating that PT systems are capable of producing a highly diverse array of terpenoids.

Family 2-associated TCs can be classified into two groups: 2-MIBS-like cyclases, and geosmin synthase (GS)-like cyclases (**Fig. S6**). 2-MIB is a monoterpenoid derivative that is formed from the cyclization of 2-methylgeranyl diphosphate. Geosmin is a diterpenoid resulting from the cyclization of farnesyl diphosphate. Structurally, the 2-MIBS-like cyclases contain a single TC domain near the C-terminal half of the protein while the first 100 to 120 amino acids are usually unannotated and often predicted to be disordered (**Fig. 4C** and **Fig. S4**). In contrast, GS-like cyclases contain two TC domains. Sequence alignments show that most 2-MIBS-like cyclases contain a conserved glycine, proline, and alanine-rich region within the unannotated N-terminal domain (consensus: GPTGLGT) (**Fig. S4**). Similarly, GS-like cyclases contain a conserved GPTGLGTSAAR (consensus) sequence between the two cyclase domains which is repeated at the very C-terminus of the protein (**Fig. S4**). These conserved motifs located in unannotated and disordered regions of TCs may function as targeting sequences responsible for mediating cargo encapsulation.

Family 2B-associated PTs are highly diverse and likely capable of producing linear isoprenoids of varying lengths (**Fig. S7**). Similar to other Family 2B cargos, encapsulin-associated PTs have a disordered N-terminal domain of 50 to 100 residues (**Fig. 4C** and **Fig. S4**). No conserved sequence motifs that may function as targeting tags could be identified. However, the consistent presence of unannotated and disordered domains may suggest their involvement in PT cargo encapsulation.

124 Family 2B systems exclusively found in *Streptomyces* contained enriched *xylB* genes encoding for XKs (**Fig. 4**). All XK gene clusters contained two distinct Family 2B encapsulins indicating that the formation of a putative two-component shell might be essential for these systems. In contrast to the other identified Family 2 cargo types, XKs do not consistently contain stretches of predicted disorder or unannotated domains. Commonly enriched accessory components such as acetylxylan esterases, xylose repressors (*xylR*), and xylose isomerases (*xylA*) suggest that these Family 2B systems may be involved in xylose utilization and metabolism (**Fig. S5**).

### Family 3 – Natural Product Encapsulins

We identified 132 Family 3 encapsulins encoded in a variety of different natural product biosynthetic gene clusters (BGCs) (**Fig. 5**). 97 Family 3 encapsulins can be found in Actinobacteria, 34 in Proteobacteria, and one in Chloroflexi. We categorized Family 3 encapsulins according to their sequence similarity and surrounding BGC type into 6 classes (**Fig. 5B**). Classes were named based on the most prominent genera encoding a given class. Family 3 BGCs encode diverse components but commonly found genes include sulfotransferases, short-chain dehydrogenases (SDRs), polyketide synthases (PKSs), non-ribosomal peptide synthetases (NRPSs), and amino-group carrier proteins (AmCPs).

**Fig. 5.**
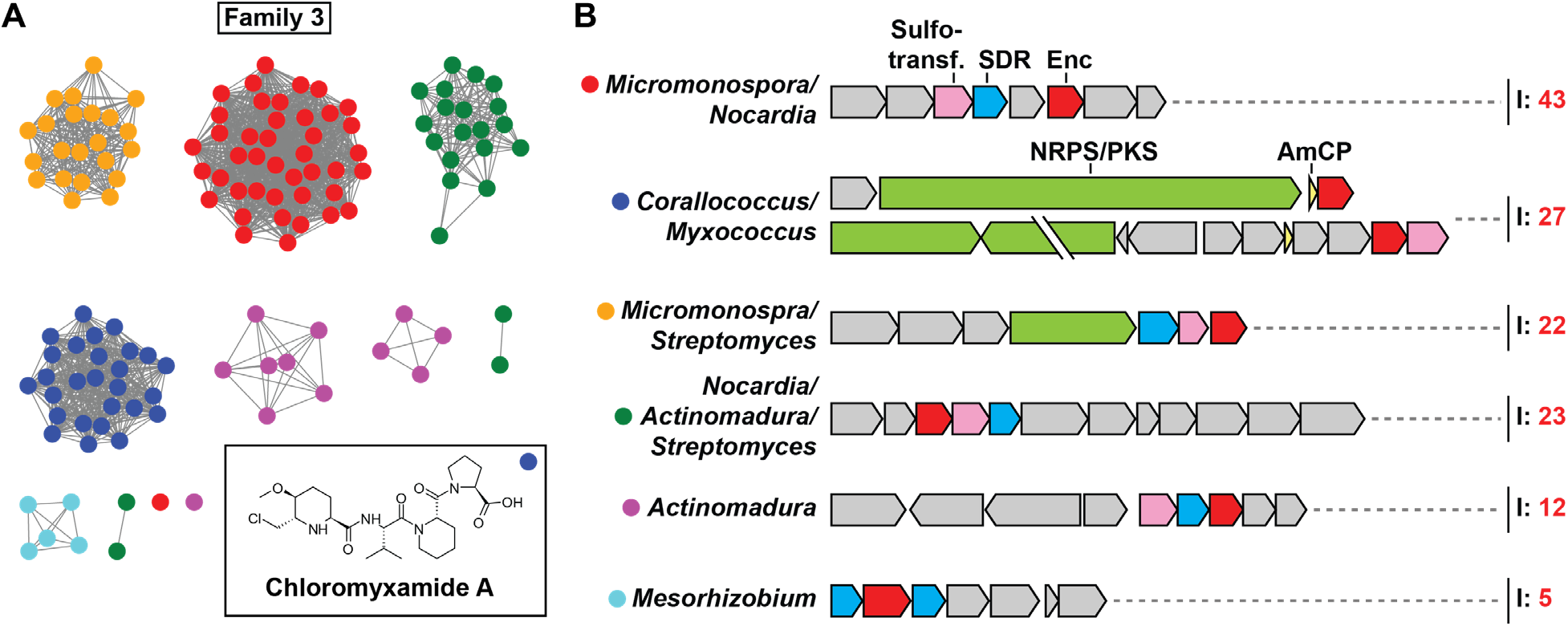
Overview of Family 3 encapsulin systems. A) SSN of Family 3 containing 138 nodes representing encapsulin capsid sequences clustered at 55% sequence identity. The inset shows chloromyxamide A, a natural product produced by a biosynthetic gene cluster encoding a Family 3 encapsulin found in *Myxococcus* sp. MCy10608.^39^ B) Diversity of operon types encoding Family 3 encapsulins. Sulfotransferases, SDR-family oxidoreductases, non-ribosomal peptide synthetases (NRPSs)/polyketide synthases (PKSs), and amino-group carrier proteins (AmCPs) are commonly found in Family 3 operons. I: number of identified operons.

Only one Family 3 encapsulin-containing BGC has been studied experimentally, namely, a system found in *Myxococcus* sp. MCy10608 (**Fig. S8**).^39^ This *Myxococcus* BGC was shown to produce a variety of chlorinated 6-chloromethyl-5-methoxypipecolic acid-containing peptide natural products dubbed chloromyxamides. The chloromyxamide biosynthetic pathway and the role of the BGC-encoded Family 3 encapsulin are currently unknown. Based on gene annotations, putative biosynthetic pathways for all the other BGC classes have been proposed (**Fig. S9, S10, S11** and **S12**). Given the presence of conserved pairs of sulfotransferases and SDRs in many of the identified BGCs, it is likely that the respective natural products will contain sulfated hydroxyl groups generated through the successive action of SDRs and sulfotransferases (**Fig. S9**).^40^ Some of the identified BGCs encode LysW-like AmCPs, suggesting that these biosynthetic pathways rely on covalently tethered intermediates, as observed in bacterial lysine and arginine biosynthesis (**Fig. S10**).^41^ Other BGCs contain large genes encoding NRPS or PKS multidomain enzymes responsible for non-ribosomal peptide and polyketide assembly (**Fig. S11**). The diversity of peptide bond-forming as well as peptide tailoring enzymes encoded in Family 3-associated BGCs suggests that they are capable of producing a structurally diverse set of peptide natural products.

A unique type of Family 3 encapsulin containing a C-terminal extension annotated as a major facilitator superfamily (MFS) domain – containing 4 to 5 predicted transmembrane helices – was identified in a number of *Mesorhizobium* spp. (**Fig. S12** and **Fig. S13**). The respective BGCs further encode SDRs and enzymes commonly found in serine biosynthesis like phosphoserine aminotransferase, and phosphoserine phosphatase. These *Mesorhizobium* BGCs might be involved in the biosynthesis of a phosphorylated amino acid derivative (**Fig. S12**). While the role of the predicted transmembrane helices found in these Family 3 encapsulins is unknown, they may form a hydrophobic MFS-like gated channel surrounding the encapsulin pores (**Fig. S13**). Alternatively, they may mediate encapsulin-lipid membrane interactions or even recruit a lipid layer around the Family 3 encapsulin shell, similar to a viral envelope.

What role do Family 3 encapsulins play in the identified BGCs? Many of the tailoring enzymes found in Family 3 BGCs contain extended unannotated or possibly disordered regions at their N- or C-termini. This may suggest that some of them are cargo proteins that are actively encapsulated, a theme observed for both Family 1 and Family 2 systems. Active encapsulation of certain biosynthetic enzymes may allow Family 3 encapsulins to function as nanoscale reaction vessels and sequester reactive aldehyde or ketone intermediates (**Fig. S8, S9, S10, S11** and **S12**) thus preventing potentially toxic side reactions in the cell cytoplasm. Similar molecular logic has been observed for bacterial microcompartments where a protein shell acts as a diffusion barrier for volatile or reactive pathway intermediates.^42^

### Family 4 – A-domain Encapsulins

A-domain Encapsulins are the most distinctive type of encapsulin-like system discovered in this study. They represent a highly truncated version of the HK97-fold and are predominantly found in genomes of hyperthermophilic Archaea. All so far sequenced *Pyrococcus* and *Thermococcus* genomes contain at least one but oftentimes two A-domain Encapsulin systems. Outside Archaea, A-domain Encapsulins are only present in the two thermophilic Bacteroidetes genera *Rubricoccus* and *Rhodothermus* (**Fig. S14**). The fact that all organisms encoding Family 4 encapsulins are thermophilic anaerobes and were all isolated from submarine hydrothermal vents may implicate these systems in biological functions directly related to the extreme environmental conditions of these unique habitats.

SSN-analysis of all 95 identified Family 4 encapsulins revealed clear separation into 3 distinct clusters from now on referred to as Family 4A, 4B and 4C (**Fig. 6A**). Four conserved operon types could be identified based on the identity of the enzymatic components encoded upstream of the A-domain protein (**Fig. 6B**). Further, a subset of identified A-domain encapsulins, including all bacterial representatives, did not have any clearly associated enzymatic components and are thus referred to as Orphan/Unknown. However, it should be noted that heme biosynthesis components are enriched in the genome neighborhood of bacterial A-domain Encapsulins (**Fig. S14**). The four conserved enzymatic components found in archaeal systems are: [NiFe] sulfhydrogenase (Hydrogenase, four subunits: αβγδ), osmotically inducible protein C (OsmC), glyceraldehyde-3-phosphate dehydrogenase (GAPDH) and deoxyribose-phosphate aldolase (DeoC). Mapping these four operon types onto the SSN showed that Hydrogenase and OsmC operons are confined to Family 4A while GAPDH and DeoC operons can only be found in Family 4B (**Fig. 6A**). Generally, each *Pyrococcus* and *Thermococcus* species encodes two separate A-domain Encapsulin systems, specifically, one Family 4A and one Family 4B operon.

**Fig. 6.**
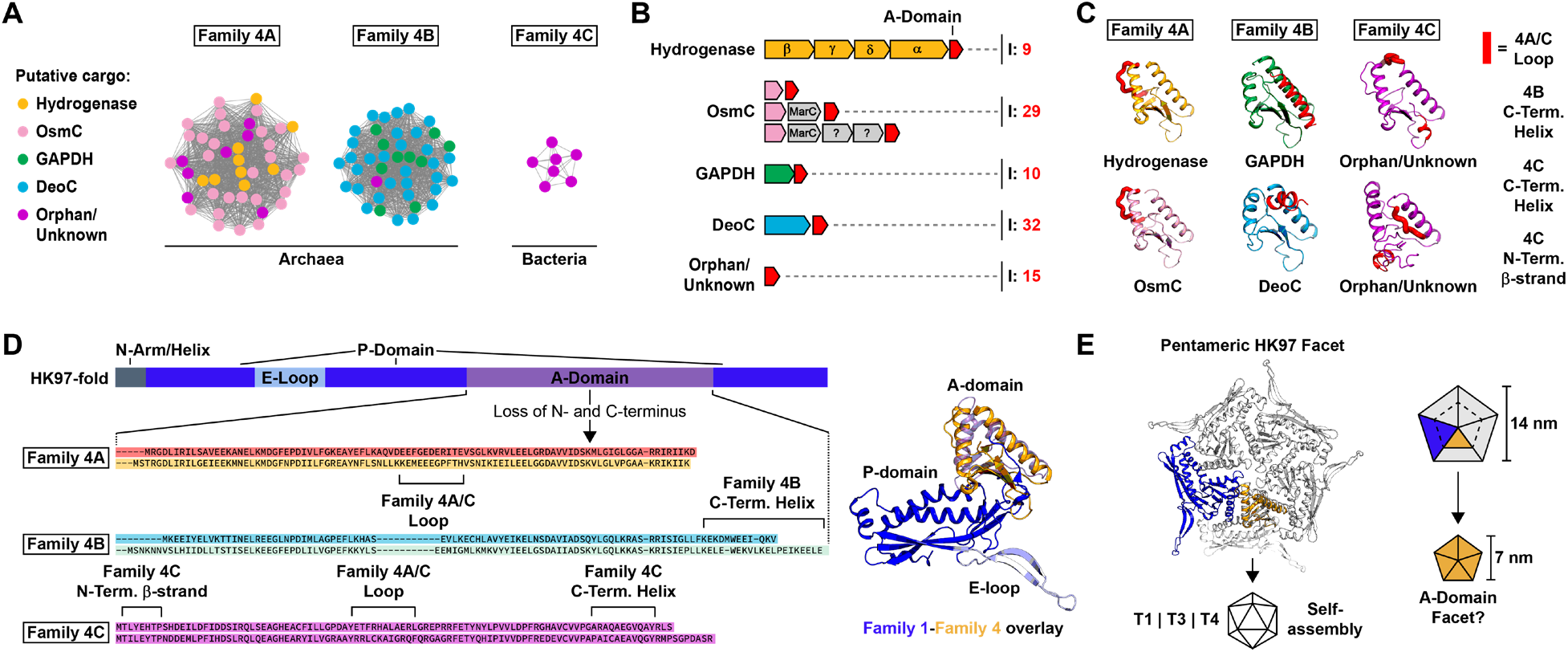
Overview and analysis of Family 4 encapsulins. A) SSN analysis of Family 4. Nodes represent 95 A-domain Encapsulins clustered at 38% sequence identity. Nodes are colored by operon type based on associated enzyme components. B) Overview of Family 4 operon types highlighting enzyme components (colored) and the number of identified systems (I). C) Structural analysis of A-domain Encapsulin monomers based on homology modelling. Structural features distinguishing Family 4A, 4B and 4C are highlighted in red. D) Left: Sequence and structure of the HK97-fold and origin of A-domain Encapsulins. Loss of N- and C-terminal domains results in a truncated protein corresponding to the A-domain of the HK97-fold. Distinguishing structural features of Family 4A, 4B and 4C shown in C) are highlighted. Right: Structural comparison of HK97-fold (T1 Classical Encapsulin (3DKT), blue and purple) and A-domain Encapsulin (yellow). E) Pentameric facet of a Family 1 encapsulin compared with an A-domain Encapsulin. A-domain Encapsulins may assemble into smaller pentameric facets about half the size of HK97-fold facets.

A-domain Encapsulins are structurally similar to the A-domain of the HK97-fold. The crystal structure of a Family 4A Hydrogenase A-domain Encapsulin from *Pyrococcus furiosus* was solved, but not further characterized (PDB ID: 2PK8).^43^ The protein was N-terminally His-tagged and crystallized as a dimer. Using 2PK8 as a threading template, the I-TASSER server^44^ was used to generate homology models of A-domain Encapsulins from Family 4A, 4B and 4C as well as all operon types (Hydrogenase, OsmC, GAPDH and DeoC) (**Fig. 6C**). Similar to the A-domain of the HK97-fold,^25,26^ A-domain Encapsulin monomers consist of two α-helices surrounding a central four-stranded β-sheet called the β-hinge. The 3 subfamilies differ due to the presence of an N-terminal α-helix or an additional C-terminal β-strand as well as the presence or absence of an extended loop between the two main helices. Sequence similarity between A-domain Encapsulins and HK97-fold proteins is very low, however, based on structural alignments (**Fig. 6D**), it appears that large portions of the HK97-fold N- and C-terminal domains were lost, resulting in a contiguous stretch of about 100 amino acids representing the A-domain. All known HK97-fold proteins have the ability to self-assemble into pentameric C5 symmetrical complexes, also known as facets, that usually assemble further into icosahedral closed capsids (**Fig. 6E**).^25,26^ HK97-fold A-domains are also crucial for the formation of symmetrical pores at the 5-fold symmetry axis in both Classical Encapsulins and viruses.^25,26^ The two main helices of the A-domain form the major interaction interfaces between the five subunits of a facet. The conformational similarity of A-domain Encapsulins and HK97-fold proteins when part of a pentameric facet can be easily illustrated via structural alignments (**Fig. 6E**). We hypothesize that A-domain Encapsulins should also be able to self-assemble into facets and potentially larger complexes. The fact that 2PK8 did crystallize as a dimer may be an artefact due to the presence of an N-terminal His-tag which could easily interfere with facet formation.

Family 4 Hydrogenase systems encode a four subunit [NiFe] hydrogenase as their enzymatic component. The specific [NiFe] hydrogenases associated with A-domain Encapsulins generally form cytoplasmic soluble heterotetrameric complexes^45,46^ and catalyze the reversible interconversion of H2 to two protons and two electrons.^47^ The A-domain Encapsulin-associated [NiFe] hydrogenase of *P. furiosus* has been partially functionally characterized, however, this was done through whole cell measurements and heterologous expression experiments which did not yield any information about the associated A-domain Encapsulin.^48,49^ In *P. furiosus*, this hydrogenase complex is known as sulfhydrogenase I (SHI) referring to its ability to act as a sulfur reductase, oxidizing H2 whilst simultaneously reducing elemental sulfur or polysulfides to hydrogen sulfide (H2S).^50,51^ SHI has been proposed to primarily work in the direction of H2 formation in an NADPH-dependent manner.^52^ It has been suggested that SHI mostly serves as a safety valve to remove excess reducing equivalents from the cytosol, thus playing an important role in maintaining intracellular redox homeostasis.^53,54^

The OsmC system encodes a single copy of the OsmC protein as its enzymatic component and often a MarC-like transmembrane protein, all located directly upstream of the A-domain Encapsulin. OsmC-type proteins are also known as organic hydroperoxide resistance (Ohr) proteins.^55^ OsmC-like proteins are known to be organic hydroperoxidases and play important roles in microbial resistance against a broad range of fatty acid hydroperoxides and peroxynitrites generated as a result of oxidative and nitrosative stress.^56^ OsmC proteins generally form dimeric structures containing a two-cysteine active site.^57-59^ Peroxides are reduced to the corresponding alcohols and water with concomitant formation of a disulfide bond between the two active site cysteines.^60^ After re-reduction, OsmC is ready for the next catalytic cycle. Studies indicate that the biological reductant of OsmC is dihydrolipoamide and not one of the more common cellular reducing agents like thioredoxin or glutathione.^61-63^ It is unclear how MarC could be involved in the function of OsmC type A-domain Encapsulin systems.^64^

The GAPDH system consists of a gene encoding for a glyceraldehyde-3-phosphate dehydrogenase arranged in a two-gene operon with the downstream A-domain component. GAPDH is a housekeeping gene present in all domains of life and is a key component of glycolysis and gluconeogenesis as well as other varied pathways and processes.^65-67^ In Archaea of the genera *Pyrococcus* and *Thermococcus*, tetrameric GAPDH is part of the reversible modified Embden-Meyerhof-Parnas (EMP) pathway responsible for glycolysis and gluconeogenesis.^68-70^ In the classical EMP pathway for sugar degradation in eukaryotes and bacteria, GAPDH catalyzes the reversible oxidation of glyceraldehyde-3-phosphate (GAP) to 1,3-bisphosphoglycerate (1,3BPG). In contrast, hyperthermophilic Archaea skip 1,3BPG formation and convert GAP directly to 3-phosphoglycerate (3PG) via enzymes only found in Archaea (GAPOR: GAP oxidoreductase or GAPN: non-phosphorylating GAP dehydrogenase).^71-75^ *Pyrococcus* and *Thermococcus* species encode GAPN which functions in the catabolic (glycolysis) direction while the single GAPDH encoded in their genomes, which is associated with an A-domain Encapsulin, is most highly expressed under gluconeogenic conditions and likely functions exclusively in the anabolic (gluconeogenesis) direction.^48,75-77^ This likely implicates GAPDH A-domain Encapsulin operons in central carbon metabolism, specifically gluconeogenesis.

DeoC systems encode a deoxyribose-phosphate aldolase upstream of the A-domain component. DeoC forms a tetrameric complex and catalyzes the reversible reaction of 2-deoxy-D-ribose 5-phosphate to GAP and acetaldehyde.^78,79^ DeoC activity facilitates the utilization of exogenous nucleosides and nucleotides for energy generation where GAP and acetaldehyde can enter glycolysis and the citric acid cycle, respectively.^80,81^ DeoC has also been shown to be upregulated under various stress conditions in *Thermococcus* species and other organisms which was hypothesized to indicate a redirection of carbon flux through DeoC and thus DNA precursor biosynthesis to maintain equilibrium between various catabolic and anabolic metabolic intermediates.^82-84^ Based on this analysis, we suggest that DeoC A-domain Encapsulin systems are involved in the utilization of nucleosides and nucleotides.

After discussing potential biological functions of A-domain Encapsulin systems, the function of the A-domain protein itself remains speculative. It is likely that A-domain Encapsulins fulfill a structural function in analogy to all other known HK97-fold proteins and that they retain the ability to self-assemble into higher order structures. We hypothesize that the respective enzymatic components of each operon type form complexes with the structural A-domain component. This is supported by a proteomics study carried out in *P. furiosus* that showed that GAPDH and the respective A-domain Encapsulin form a stable complex.^46^ Complex formation might be based on an interaction of the enzymatic component with one or multiple A-domain facets or even the encapsulation of the enzyme component inside a closed shell formed from self-assembled A-domain oligomers. What could be the specific functional role A-domain proteins play in these complexes? One possibility is that A-domain Encapsulins stabilize the respective enzymatic components through close association or encapsulation, in essence acting as specialized molecular chaperones.^85,86^ This might result in increased thermal stability, increased resistance against oxidative stress and a prolonged productive lifetime of the associated enzyme complexes. A-domain Encapsulins might also protect enzymatic reaction intermediates from competing side reactions or sequester reactive or toxic intermediates inside a protein shell similar to what has been proposed for bacterial microcompartments.^42^

### Pathogen-encoded encapsulins and their role in pathogenicity and virulence

Encapsulin systems can be found in a wide variety of prominent Gram-negative and Gram-positive pathogens. Family 1 and 2 encapsulins (peroxidase, Flp, and desulfurase systems) are found in pathogenic *Escherichia coli, Klebsiella pneumoniae*, and *Acinetobacter baumannii*, belonging to the highly virulent and antibiotic-resistant ESKAPE group of pathogens, responsible for the majority of life-threatening hospital-acquired infections worldwide.^87^ Family 1 peroxidase operons are widely distributed in Mycobacteria, including *M. tuberculosis* and *M. leprae*, the causative agents of tuberculosis and leprosy, respectively.^88,89^ Flp and desulfurase systems are both found in *Burkholderia cepacia* (pulmonary infections, cystic fibrosis) and *Burkholderia pseudomallei* (melioidosis)^90^ while *Nocardia* spp. (nocardiosis), *Bordetella* spp. (whooping cough), and *Clostridium* spp. (colitis, botulism, gangrene) encode peroxidase and Flp encapsulins.^91,92^

Most pathogen-encoded encapsulins are likely involved in stress resistance and nutrient utilization functions, often important for host invasion and proliferation in hostile environments like an infection site.^93-95^ A direct link between *M. tuberculosis* oxidative stress resistance during infection and a Family 1 peroxidase system has recently been established which represents the first direct evidence of the involvement of encapsulins in pathogenicity and virulence.^11^ In addition to stress resistance, it is possible that specialized encapsulin-based nutrient utilization systems – specifically for the two scarce and essential elements iron and sulfur – can increase pathogen fitness and proliferation, similar to the importance of bacterial microcompartment-based carbon and nitrogen source utilization systems for the pathogenicity of *Salmonella typhimurium* (food poisoning), *Enterococcus faecalis* (nosocomial infections), and *Clostridium difficile* (colitis).^96-98^ Future efforts to characterize pathogen-associated encapsulin systems may yield novel targets for therapeutic intervention.

### Phylogenetic analysis of encapsulins and related HK97-fold proteins

All four families of encapsulins discussed above belong to the Pfam clan CL0373. A Pfam clan is a collection of related Pfam families.^19^ Membership within a Pfam clan is determined by up to four independent pieces of evidence: related structure/fold, related function, significant matching of sequences to HMMs from separate Pfam families, and pairwise profile-profile HMM alignments based on HHsearch.^99^ The fact that Pfam clans are meant to contain only Pfam families that share a common evolutionary origin^19^ is a first indication that all four encapsulin families are in fact evolutionarily related to the other HK97-fold proteins, all representing phage and virus capsid proteins, contained within CL0373. To further investigate the relationship between encapsulin-like proteins and virus capsids, we carried out a detailed phylogenetic analysis of CL0373. Due to the generally low sequence similarity among virus capsid proteins and between encapsulin families which makes multiple sequence alignments difficult, we based our analysis on the most conserved regions of the HK97-fold, specifically the A-domain and neighboring regions belonging to parts of the HK97-fold P-domain (**Fig. 6D**). The resulting phylogenetic tree showed relatively confident bootstrap values and allowed us to investigate the relationships between all members of CL0373 in more detail (**Fig. 7A**). All encapsulin families except Family 4 (A-domain Encapsulins) are more closely related to one another than to other HK97-type proteins indicating that they might all share a recent common ancestor. The P22 coat protein family (PF11651) seems to be the virus capsid protein family most closely related to Family 1, 2 and 3 encapsulins. Classical Encapsulins (Family 1) found in both Bacteria and Archaea are more closely related to one another than to any other HK97-fold proteins. This suggests inter-domain horizontal transfer of Family 1 encapsulin systems, likely from Bacteria to Archaea, which is a well-documented phenomenon.^100,101^ A-domain Encapsulins (Family 4) appear to be more evolutionarily distinct from the other encapsulin families and to be more closely related to other HK97-fold capsid proteins. The Gp23 family (PF07068) generally found in T4-like bacteriophages seems to be the most distantly related Pfam family compared with Family 1, 2 and 3 encapsulins. Our sequence-based analysis suggests that encapsulins share common ancestry with all HK97-fold families contained within CL0373 and indicates that encapsulin systems likely evolved from viruses, specifically from members of the widespread virus order Caudovirales.^102^

**Fig. 7.**
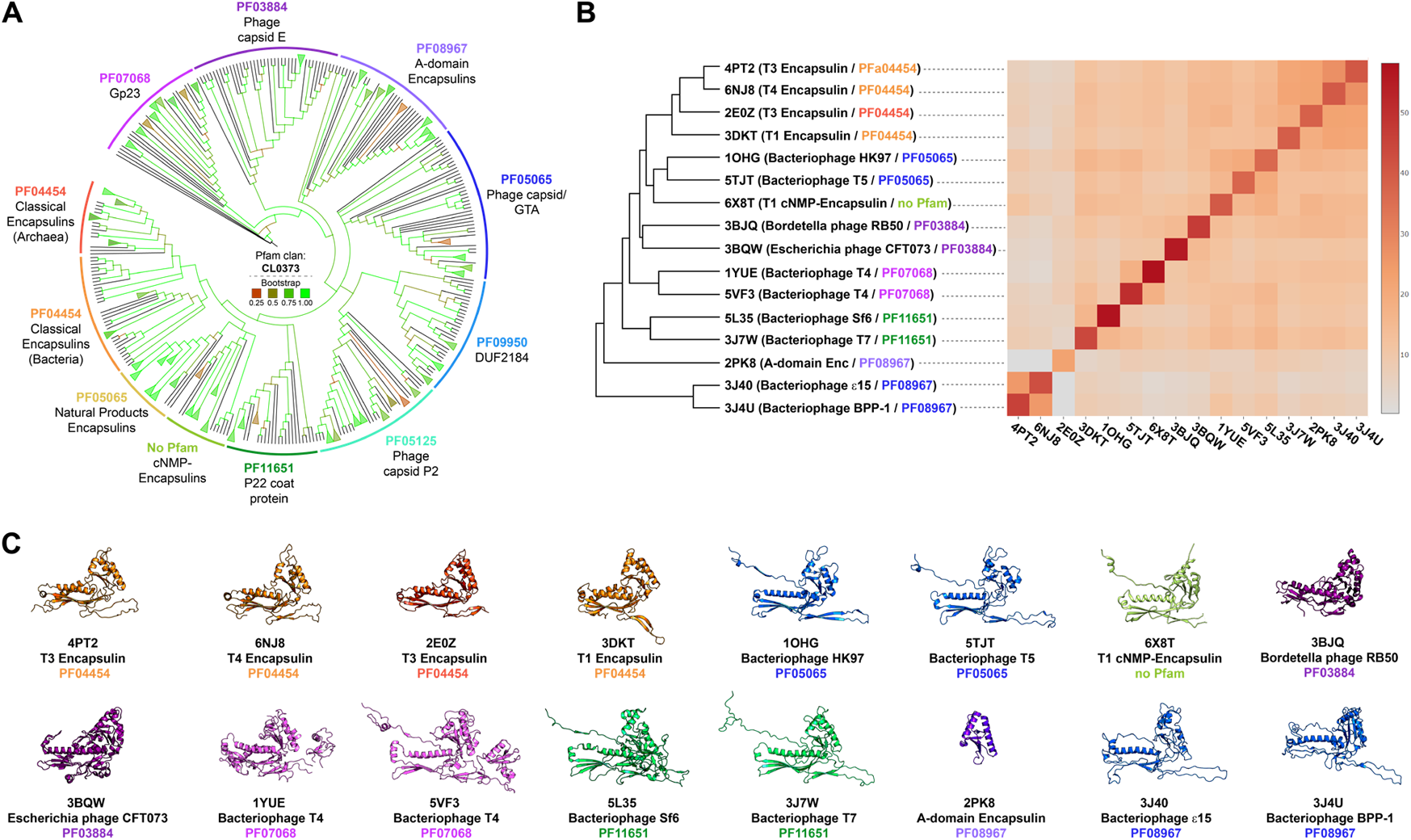
Phylogenetic analysis of HK97-fold proteins. A) Phylogenetic tree of Pfam clan CL0373. Branches colored by Bootstrap values. B) DALI structural comparisons of representative CL0373 structures of each family with available structures. Left: dendrogram based on pairwise Z score comparisons. Right: matrix/heatmap representation of Z scores based on pairwise comparisons. The color scale indicates Z scores. Pfam families colored as in A). C) Representative monomer structures used in B) for structural comparisons colored by Pfam family color. PDB IDs, names and Pfam families shown below each monomer structure.

Further analysis of CL0373 members was carried out via structural comparisons of representatives of all investigated Pfam families for which structures were available (**Fig. 7B**). Pairwise structural similarities were evaluated using the DALI Z score.^103^ The Z score is a measure of the overall quality of a given structural comparison. All-against-all structural comparisons showed that Family 1 encapsulins form an apparent monophyletic cluster while the single available Family 2 encapsulin (6×8T)^8^ was more similar to PF05065. It should be noted that 6×8T represents a Family 2 encapsulin without a cNMP-binding domain. No structure of the more ubiquitous cNMP-containing Family 2 systems is currently available. The other viral HK97-type proteins are more divergent compared to the available encapsulin structures (**Fig. 7B**). Further visual inspection of sample structures (**Fig. 7C**) reveals that Family 1 encapsulins do not possess an extended N-arm which is present in the majority of other HK97-fold proteins. Instead, they possess an N-terminal helix which forms part of the binding pocket for the targeting peptide of cargo proteins.^10,15^ Some of the sample structures additionally possess insertion domains that are present in the E-loop (PF07068).^25,26^ Family 1 encapsulins generally appear more compact with a shorter central P-domain helix and shorter E-loop. In accordance with the DALI structural comparison, the Family 2 encapsulin example appears structurally more similar to the phage capsid family PF05065 than to Family 1 encapsulins. As discussed above, Family 4 encapsulins are structurally similar to the A-domain of the HK97-fold.^43^ No Family 3 structures have been solved at the time of writing. However, some Family 3 members have been annotated as belonging to Pfam family PF05065 which may indicate that they are more structurally similar to Family 2 than Family 1 encapsulins.

Both our sequence- and structure-based analysis argues for a viral origin of encapsulin systems likely via domestication of prophage HK97-type capsid proteins. This is also in agreement with the fact that HK97-fold viruses are ubiquitous and found as proviruses and prophages in the genomes of members of all domains of life while encapsulins show a narrower distribution.^9,102^ Considering that one of the current hypotheses regarding the origin of viral capsid proteins is a scenario where they ultimately derive from cellular protein folds of ancient or extinct cellular lineages,^104^ the HK97-fold might have undergone a re-recruitment, and as part of encapsulin systems has now returned to its cellular origin.

## Conclusion

The curated set of encapsulin-like systems discovered and analyzed here, sheds light on the true functional diversity of microbial protein compartments. Proposed encapsulin functions include roles as reaction spaces for various anabolic (Family 2 and 3) and catabolic (Family 2) processes, storage compartments (Family 1), enzyme regulatory systems (Family 2 and 4) as well as chaperones (Family 4). Encapsulins are found in aerobic and anaerobic microbes that occupy nearly all terrestrial and aquatic habitats as well as host-associated niches. Additionally, encapsulins are widespread in bacterial and archaeal extremophiles, specifically (hyper)thermophiles and acidophiles. The evolutionary scenario outline above, where encapsulin systems are the result of the molecular domestication of phage capsid proteins by cellular hosts, is further supported by the existence of transitional systems like the Family 1 encapsulin found in *Sulfolobus solfataricus* whose genetic context indicates that it used to be part of a now defective prophage.^36^ It is possible that other viral capsid protein folds may also have undergone a similar recruitment process and now serve specific host metabolic functions. This idea is supported by the recent description of the involvement of the retrovirus-like capsid protein Arc in inter-neuron nucleic acid transport.^105^ In conclusion, our study establishes encapsulins as a ubiquitous and diverse class of protein compartmentalization systems and lays the groundwork for future experimental studies aimed at better understanding the physiological roles and biomedical relevance of encapsulins.

## Methods

### Genome-mining searches for encapsulin-like systems

Family 1 and Family 4 encapsulins were identified using the Enzyme Function Initiative-Enzyme Similarity Tool (EFI-EST) *Families* search function against the full UniProt database to filter for sequences corresponding to Pfam families PF04454 (Family 1) or PF08957 (Family 4) in May 2020.^20,106-109^ Family 4C encapsulins were identified through additional blastp searches against the NCBI_nr database using the initially identified Family 4A and 4B hits as queries. Family 2 encapsulins were initially identified based on using the EFI-EST *Sequence BLAST* function with a previously identified encapsulin as a query (WP_011055154.1).^8^ Searches were carried out against the UniProt database with an E-value of 5. This allowed us to generate an initial SSN of 1,770 sequences. To expand the dataset, we aligned 40 edge sequences from the initial dataset containing both Family 2A and 2B encapsulins using Clustal Omega v1.2.2 in the Geneious Prime software package with fast clustering (mBed algorithm) and a cluster size of 100 for mBed guide trees. Sequences were truncated to only contain the C-terminal capsid component – removing the putative cNMP-binding domain – and used to generate an initial HMM model using the hmmbuild function of the HMMER3 software package.^110,111^ This HMM was then used as an input for the HMMER search tool in the MPI Bioinformatics Toolkit (https://toolkit.tuebingen.mpg.de/). Searches were carried out against the UniProt_Trembl database in May 2020 using an E-value cutoff of 10, 0 MSA enrichment iterations in HHblits, and a maximum of 10,000 target hits.^110,112,113^ Family 3 encapsulins were identified by searching a previously identified putative encapsulin from *Myxococcus* (UniProt ID: A0A346D7L6)^39^ against the UniProt database using the EFI-EST *Sequence BLAST* function with an E-value threshold of 1 in February 2021. The resulting datasets generated from these initial searches contained the following numbers of sequences: Family 1: 2,540, Family 2: 3,859, Family 3: 215, Family 4: 95. Sequences labelled as fragments and unclassified sequences with superkingdoms labelled as metagenome were excluded. Family 1, 2, and 3 datasets were significantly contaminated with bacteriophage capsid proteins. To remove phage contamination in the Family 1 and 2 datasets, custom Blast databases were generated containing proteins encoded within 10 kb upstream and downstream of each identified capsid gene. The custom Blast databases were then searched against proteomes of HK97-type phages and a broad dataset of prokaryotic dsDNA viruses (proteome IDs: UP000002576 and UP000391682) using blastp with default settings with an E-value threshold of 0.1. Proteins identified as phage-related were excluded from the datasets. Because the Family 3 encapsulin dataset was much smaller than Family 1 and Family 2, phage proteins could be easily filtered manually by removing genome neighborhoods containing phage-associated Pfam domains (PF0860, PF03354, PF04586, PF00589, PF05135). All datasets were then further manually curated to exclude any remaining genome neighborhoods containing phage-related proteins. The final curated datasets contained the following number of sequences for each family: Family 1: 2,383, Family 2: 3,523, Family 3: 132, Family 4: 95 (**Supplementary Data 1**).

### Phylogenetic analyses and construction of phylogenetic trees

#### Encapsulin distribution in prokaryotic phyla

An initial diagram of the phylogenetic distribution of prokaryotes was constructed from a previously published maximum likelihood tree of ribosomal protein alignments using the iTOL server.^21,114^ Branches corresponding to Eukaryotes were removed, display mode set to circular, and clades were collapsed to a threshold of < 0.65 BRL. Branches were then annotated manually to highlight encapsulin containing phyla.

#### Encapsulins and related HK97-fold proteins

To infer phylogenetic relationships between encapsulin-like proteins and other HK97-type proteins, the *Phage-coat* Pfam clan CL0373 was used as a starting point.^19^ Sequences from all families found within CL0373 that contained more than 10 members were used. The following Pfam families were considered with the number of sequences used shown in parentheses: *DUF1884/Family 4 encapsulins* PF08967 (40), *DUF2184* PF09950 (37), *Gp23* PF07068 (40), *Linocin_M18/Family 1 encapsulins* PF04454 (68), *P22_CoatProtein* PF11651 (40), *Phage_cap_E* PF03864 (40), *Phage_cap_P2* PF05125 (40) and *Phage_capsid* PF05065 (40). Sequences were selected from the Seed and Full alignments of each Pfam family. Sequences from the following protein families that had no Pfam designation were additionally included in the analysis: Family 2 encapsulins (40) and Family 3 encapsulins (40). Sequences of putative Gene Transfer Agents belonging to family PF05065 were also included (29).^115^ Alignments, sequence curation and phylogenetic inference analyses were carried out using the NGPhylogeny.fr server.^116^ A custom workflow using the following tools and parameters was used. For multiple sequence alignment, MAFFT^117^ was utilized with standard parameters; for alignment curation, BMGE^118^ was used with a maximum entropy threshold of 0.75 and otherwise standard parameters; for tree inference, PhyML+SMS^119^ was used with standard parameters; for tree visualization, iTOL^114^ was used with the following parameters deviating from the pre-set: display mode: circular, branch lengths: ignore, bootstraps: display as color with range 0 to 1, auto collapse clades: BRL < 0.5. The sequence most distant to Family 1 encapsulins was used as the outgroup: J7HY26 (PF07068).

#### Terpene cyclases and polyprenyl transferases

To analyze the evolutionary relationships and diversity of terpenoid-related enzymes identified in Family 2 encapsulin operons, separate multiple sequence alignments and phylogenetic inference analyses were carried out for 530 terpene cyclase (all newly identified) and 122 polyprenyl transferase (97 newly identified, 25 already experimentally characterized)^120^ sequences (**Supplementary Data 1**). Already characterized polyprenyl transferase sequences were incorporated into our analysis to infer the putative substrate range of newly identified sequences. A custom workflow on the NGPhylogeny.fr server for sequence alignments, curation, and phylogenetic inference was used. MAFFT was utilized for multiple sequence alignments with standard parameters; alignment curation was done via BMGE and standard parameters; for tree inference, PhyML+SMS was employed using standard parameters; for phylogenetic tree visualization, iTOL was used with the following non-standard parameters: display mode: unrooted, branch lengths: ignore, bootstraps: display as color with range 0 to 1.

### Sequence similarity network analysis

Sequence similarity networks (SSNs) were calculated using the EFI-ESI server.^20,106,121^ Initial SSNs were generated for each family with edge E-values of 5 and alignment thresholds corresponding to approximately 40% sequence identity. SSNs were visualized in Cytoscape v3.8^122^ using the yFiles organic layout and were then filtered to the following percent identity thresholds to optimize cluster separation and visual presentation: Family 1: 49%, Family 2: 70%, Family 3: 55%, Family 4: 38% (**Supplementary Data 2**). Nodes were colored according to cargo type for Family 1, 2, and 4 encapsulins. Family 3 encapsulin nodes were colored according to natural product gene cluster type.

### Genome neighborhood analysis

Genome neighborhood analysis was performed using the EFI-GNT server with EFI-ESI-generated and Cytoscape-curated network files (xgmml format) as inputs resulting in computed genome neighborhoods extending 20 open reading frames up- and downstream of the identified encapsulin-encoding genes.^20,106,121^

To identify Family 1 cargo proteins, we first generated a custom database of all proteins encoded within 5000 bp up- and downstream of identified Family 1 encapsulin genes and used blastp to search for the Family 1 targeting peptide consensus sequence (SDGSLGIGSLKRS).^9^ Blastp parameters were automatically adjusted for short input sequences and an E-value of 200,000 was used. HMM templates representative of each cargo class identified through initial blastp searches were then generated and used as inputs for HMMsearch. The resulting set of cargo hits was then classified based on their Pfam or Interpro annotation. If no Pfam or Interpro annotation was present, cargo proteins were annotated based on sequence similarity. Identified cargo proteins that did not have corresponding NCBI or UniProt accession codes were labelled as *putative*. Manually curated cargo proteins that were not identified by any of the above search methods but located immediately adjacent to an encapsulin gene in the GNN or in the NCBI Nucleotide graphic interface were labelled as *manually curated*.

Family 2 cargo proteins were identified by constructing a custom database of all proteins encoded within 20 open reading frames of the identified Family 2 encapsulins, then HMMs were constructed from representative terpene cyclases, polyprenyl transferases, cysteine desulfurases, and xylulose kinases using HHbuild.^110^ The resulting HMMs were then used to query our custom Family 2 database via HMMsearch. Identified cargo proteins of encapsulins not present in the ENA database were curated manually.

Family 3 and Family 4 encapsulins were manually inspected for putative cargo proteins and operon similarity using EFI-GNT-generated genome neighborhood diagrams.

### Protein homology models and protein structure analysis

#### General

General protein structure editing and visualization was done using UCSF Chimera,^123^ UCSF ChimeraX^124^ and PyMOL.

#### Homology models

All protein homology models used for classification and analysis of Family 4 encapsulins were generated using the I-TASSER Protein Structure & Function Prediction server^44^ with standard parameters. The following sequences were used as inputs: Family 4A: F0LMI5; Family 4B: F0LIR3 and O59495; Family 4C: A0A1M6P7G0 and A0A2H0JLL0. In all cases, 2PK8 was found to be the best template.

#### Structure comparisons and similarity analysis

Structure comparisons between different HK97-type proteins were carried out on the DALI server^103^ using the following representative experimentally determined structures: PF04454: 4PT2, 6NJ8, 2E0Z and 3DKT; no Pfam (T1 Family 2A encapsulin): 6×8T; PF03884: 3BJQ and 3BQW; PF05065: 1OHG and 5TJT; PF07068: 1YUE and 5VF3; PF11651: 5L35 and 3J7W; PF08967: 2PK8; PF08967: 3J40 and 3J4U. Structural similarities between the selected proteins were evaluated based on the DALI Z score, which represents a measure of the quality of the overall structural alignment. For structure alignment visualization, structural similarity matrices resulting from all-against-all structure comparisons and the respective dendrograms were generated using the all-against-all structure comparison tool on the DALI server.

### Analysis of disordered protein sequences

Sequence disorder analyses were carried out using the Disopred3 server^125^ for the following representative proteins for each Family 2B cargo class: CD: A0A010WJT9, PT: A0A0B5EUR5, TC: 2-MIBS-like: Q9F1Y6, TC-GS-like: A0A3D0QW52. Disopred3 outputs were visualized using GraphPad Prism v9.0.2.

## Data availability

An annotated and curated spreadsheet of all identified encapsulins and cargo proteins is available as Supplementary Data 1.xlsx. Annotated SSNs for each encapsulin family (Family_1_SSN.xggml, Family_2_SSN.xggml, Family_3_SSN.xggml, and Family_4_SSN.xggml) are available as a compressed zip file (Supplementary Data 2.rar).

## Acknowledgements

We gratefully acknowledge funding from the NIH (R35GM133325).

## Author contributions

M.P.A and T.W.G designed the study, carried out computational analyses and wrote the paper.

## Supplementary information

Supplementary information containing additional data and analyses for Families 1, 2, 3, and 4 is available and contains Figs. S1-S16 and references.

