## Supplementary Information for "Large-scale computational discovery and analysis of virus-derived microbial nanocompartments"

### 1. Additional data and analysis for Family 1 Encapsulins

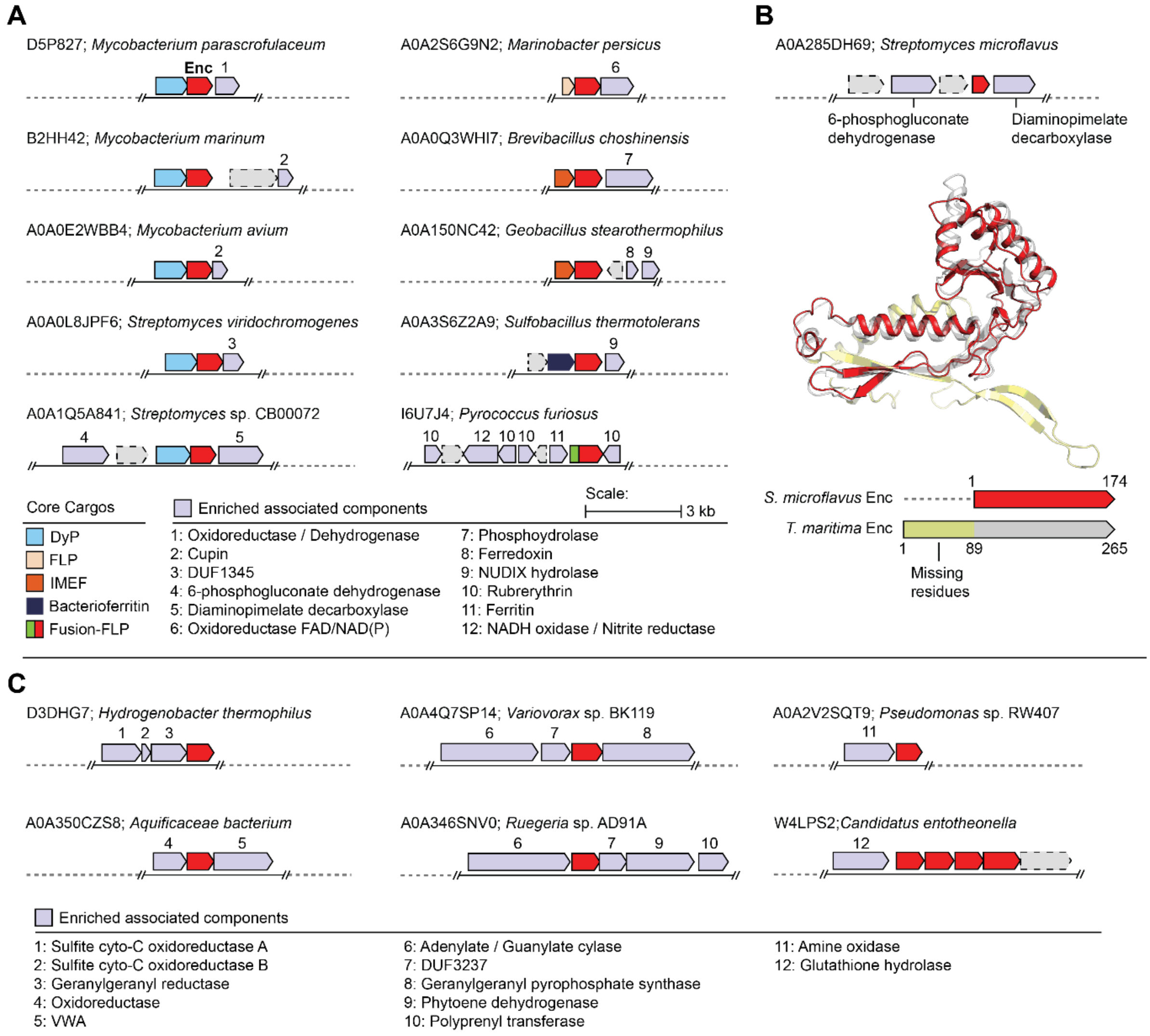

**Fig. S1.** Family 1 variants and minor operon types. **A)** Conserved operon variants containing Family 1 encapsulins (red) are illustrated for 5 of the major Family 1 cargo classes. Variably enriched components are commonly found in operons for respective cargo classes. **B)** A novel variant of short Family 1 encapsulin identified in *Streptomyces* is characterized by an N-terminal truncation that results in the loss of the N-terminal helix, E-loop, and part of the P-domain. An I-TASSER[2] model (red) was generated using a representative protein sequence (UniProt: A0A285DH69) from *Streptomyces microflavus*. Alignment with the structure of *T. maritima* encapsulin (PDB ID: 3DKT) (yellow) indicates the conserved components of the truncated encapsulin are structurally similar to other Family 1 encapsulins. **C)** Selected Family 1 operons classified as *Minor* that cannot be grouped with any of the other Family 1 cargo classes. These operons do not have clearly defined cargos or targeting peptide sequences, but often contain generally enriched components.

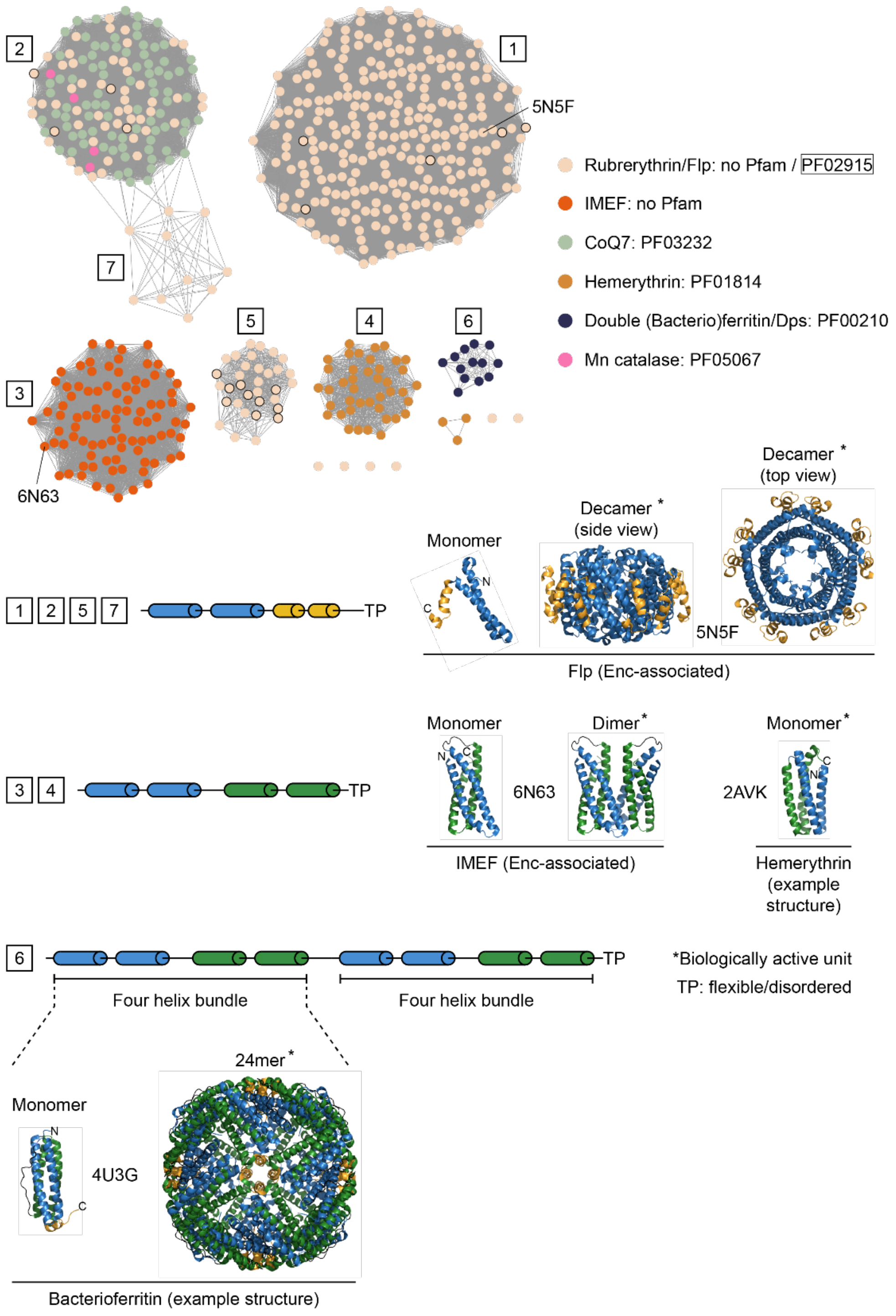

**Fig. S2.** Ferritin-like protein (FLP) superfamily cargo proteins. Top: Sequence similarity network clustered at 30% sequence identity of all identified Family 1 FLP cargo proteins. Clusters are numbered from the largest to the smallest cluster. Names and Pfam families are shown on the right. PDB IDs of structurally characterized proteins are indicated. Nodes with black outlines represent fusion Flp systems. Bottom: Secondary structures of different clusters as analysed by Jpred 4.[1] Example monomer and biologically active unit structures are shown. No encapsulin-associated hemerythrin and bacterioferritin cargo structures are available, thus, general examples are shown (PDB IDs: 2AVK and 4U3G). Cluster 6 contains unusual double four helix bundle proteins that might assemble into 12mer instead of 24mer bacterioferritin-like complexes. TP: targeting peptide.

### 2. Additional data and analysis for Family 2 Encapsulins

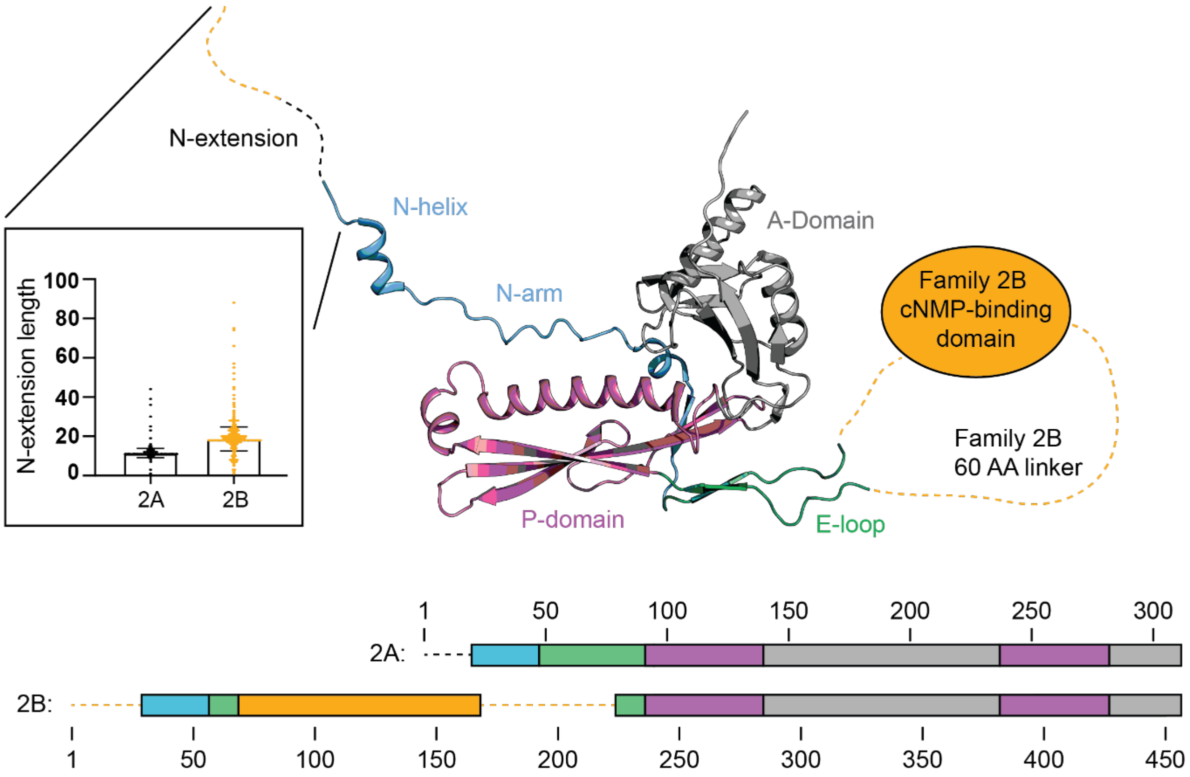

**Fig. S3.** Structural overview of Family 2 encapsulins. The structure presented here (PDB ID: 6X8M) illustrates the conserved domains for Family 2 encapsulins, namely the N-arm, E-loop, P-domain, and A-domain. Family 2B encapsulins contain a cNMP-binding domain and 60 amino acid long linker inserted within the E-loop. This feature is not present in Family 2A encapsulins. As shown by the inset graph, the N-terminal extension found in Family 2B encapsulins is generally longer than in family 2A (N-extension lengths: Family 2A: 12 ± 2 AAs; Family 2B: 19 ± 6 AAs, error bars indicate standard deviations). The inset graph was generated using GraphPad Prism v 9.0.2.

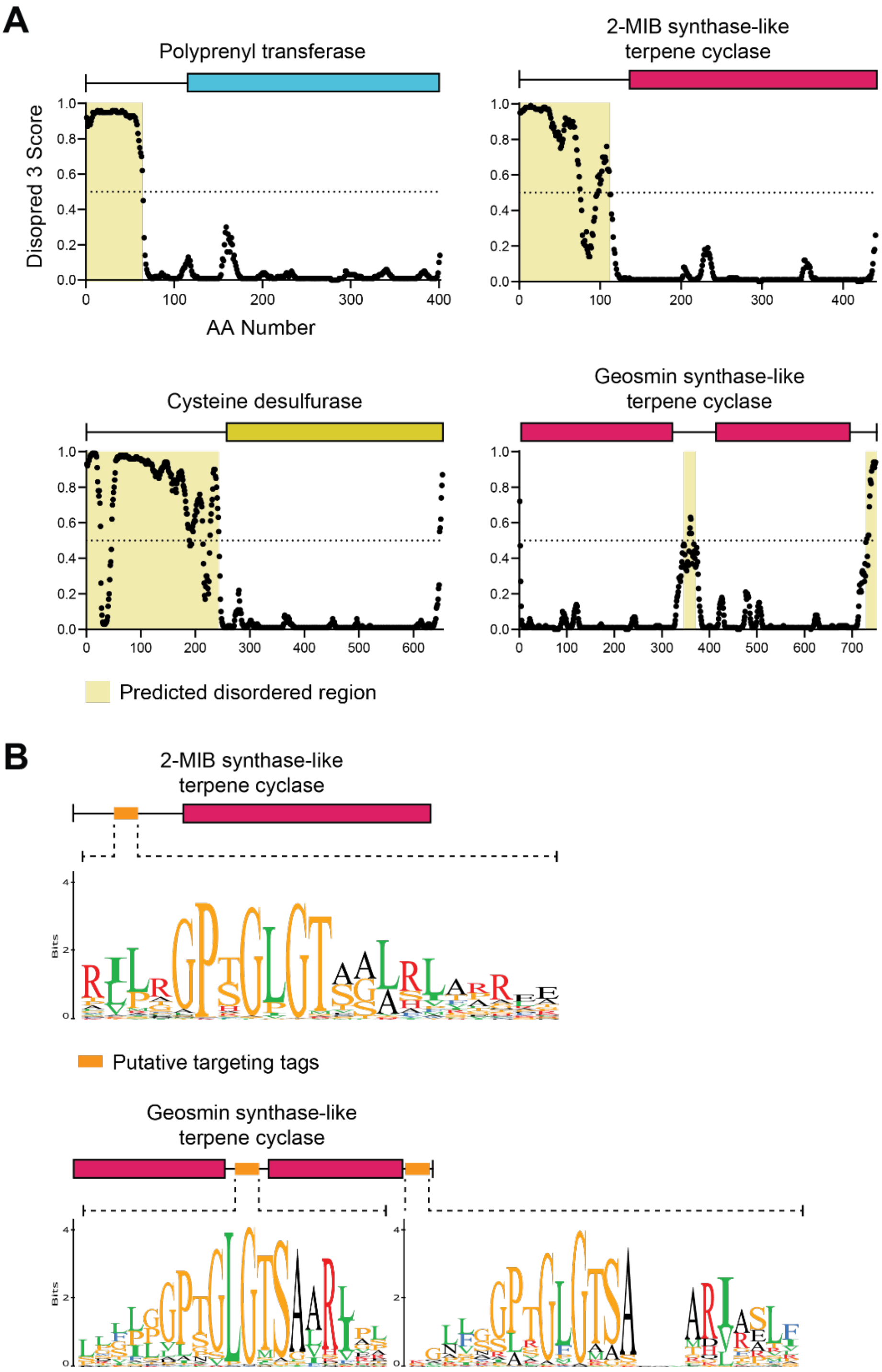

**Fig. S4.** Analysis of disordered regions in Family 2 cargo proteins. **A)** The N-termini of PTs, CDs, and 2-MIB synthase-like TCs are predicted to be disordered. Graphs illustrate per-residue disorder predictions from Disopred3[3] of representative sequences (CD: A0A010WJT9, PT: A0A0B5EUR5, TC: 2-MIB synthase-like: Q9F1Y6, GS-like: A0A3D0QW52). Residues with predicted scores of 0.5 or greater (dotted line) are designated as disordered. Regions of disorder are highlighted in yellow. Plots were created using GraphPad Prism v 9.0.2. **B)** Sequence alignments (Clustal Omega v 1.2.2 in Geneious Prime v 2020.1.2) of 499 2-MIB synthase-like TCs show a conserved N-terminal consensus sequence of GPTGLGT. Similar consensus sequences of GPTGLGTSAAR were identified within internal and C-terminal disordered regions from an alignment of 118 geosmin synthase-like TCs.

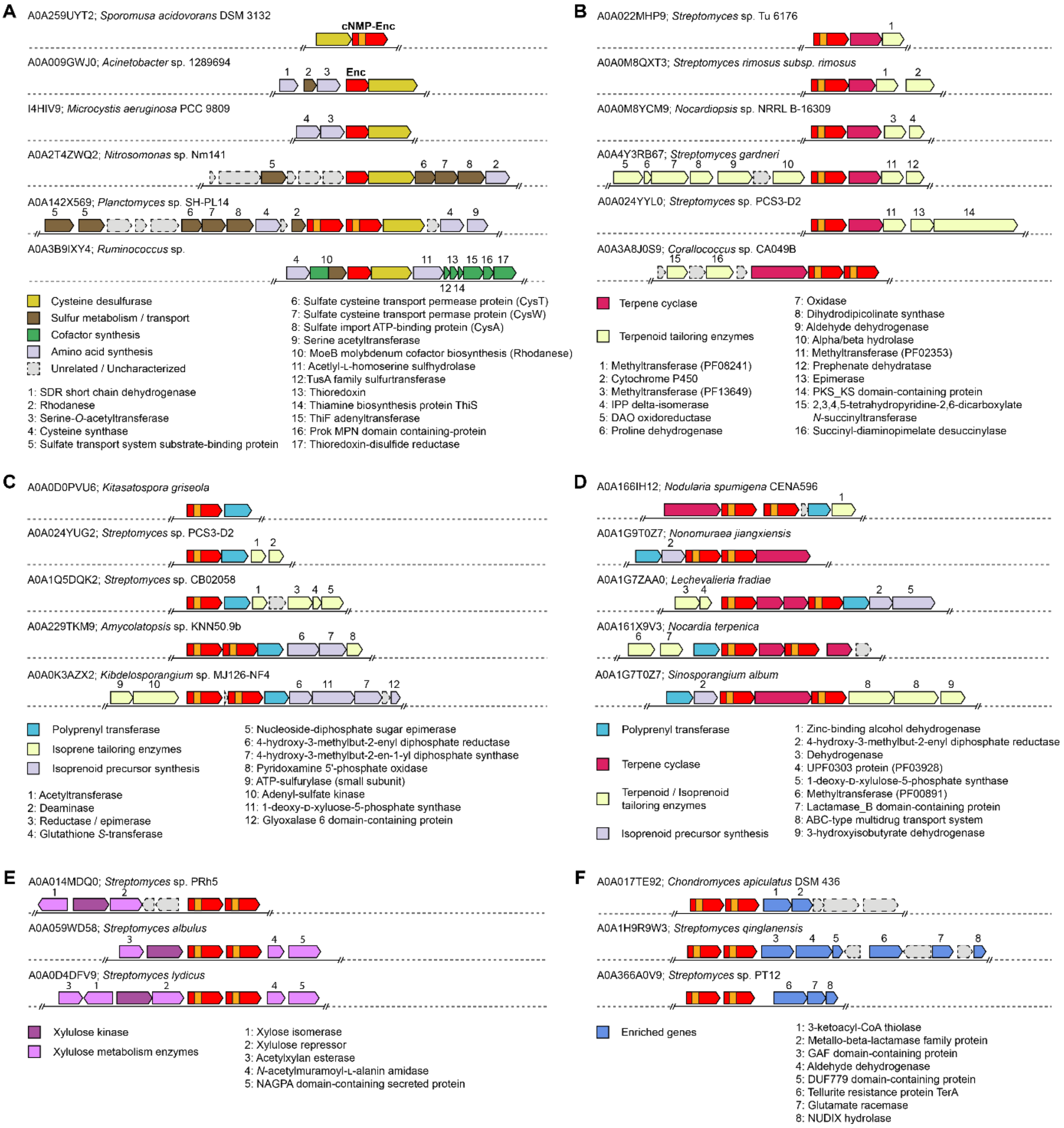

**Fig. S5.** Family 2 variants and minor operon types. **A)** Selected CD operons containing both Family 2A and Family 2B encapsulins contain enriched accessory components likely involved in sulfur metabolism and transport, cofactor synthesis, and amino acid synthesis. Serine *O*-acetyltransferases and rhodaneses are among the most commonly enriched genes in Family 2A operons. **B)** Family 2B TC operons contain various terpene tailoring enzymes. The most commonly enriched tailoring enzymes in TC systems are methyltransferases, but other enriched tailoring enzymes likely expand the diversity of terpenoids produced from these operons. **C)** Family 2B PT operons contain enriched tailoring enzymes and enzymes associated with isoprenoid precursor biosynthesis. **D)** 165 Family 2A encapsulins were found in operons containing both PTs and TCs. Similar to PT and TC operons, these mixed operons often contain enriched terpenoid and isoprenoid tailoring enzymes and isoprenoid precursor synthesis enzymes **E)** Family 2B XK operons contain highly conserved components associated with xylulose metabolism. It is worth noting that all XK operons encode for two different Family 2B encapsulins. **F)** Minor Family 2 operons containing enriched components and do not contain genes encoding for CDs, TCs, PTs, or XKs.

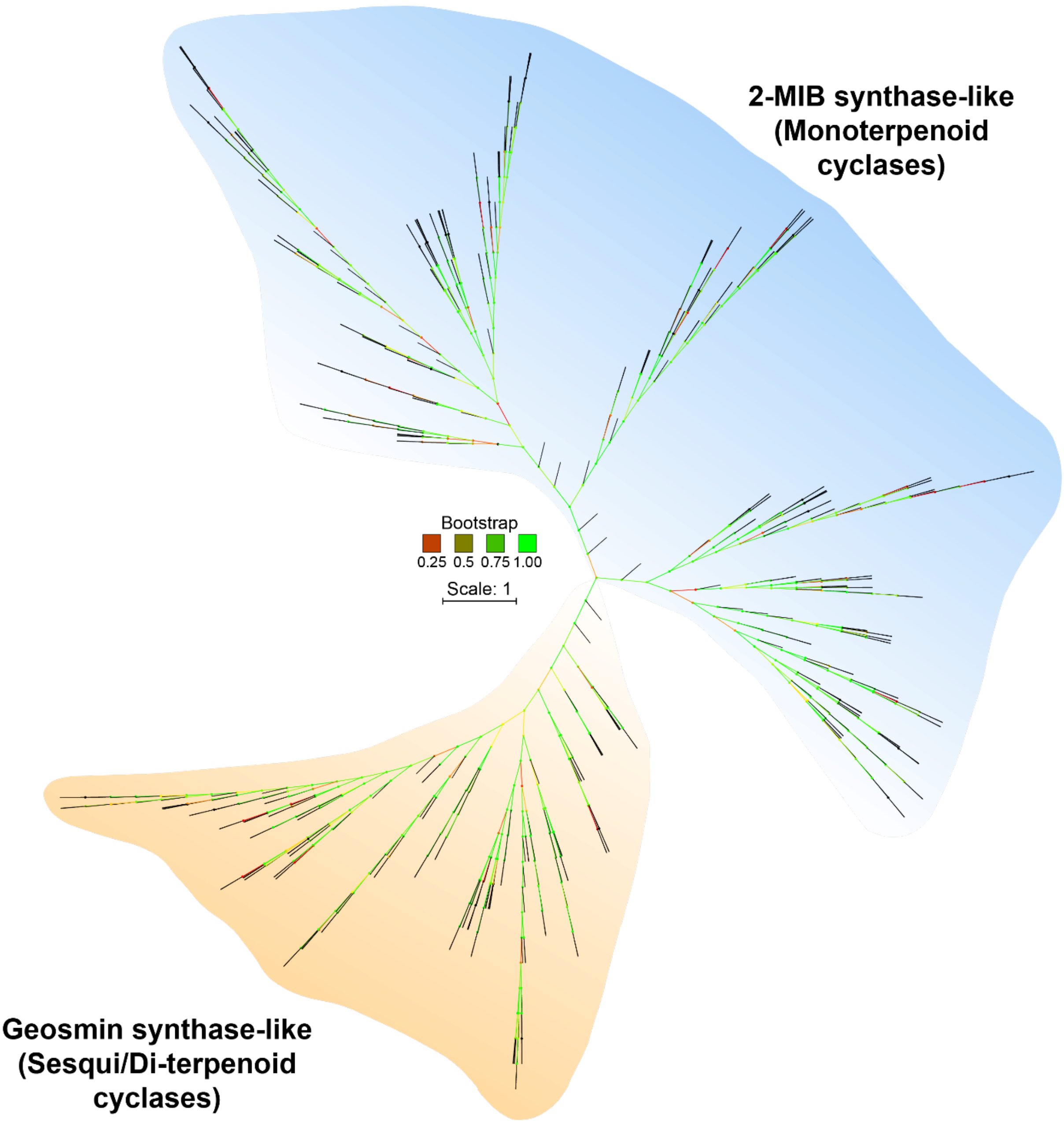

**Fig. S3.** Phylogenetic analysis of 530 Family 2-associated putative terpene cyclase (TC) cargo proteins. MAFFT was utilized for multiple sequence alignments with standard parameters; alignment curation was done via BMGE and standard parameters; for tree inference, PhyML+SMS was employed. Based on homology to characterized enzymes, Family 2-associated terpene cyclases can be divided into two major groups, one similar to geosmin synthase, likely representing sesqui- and diterpenoid cyclases, and one similar to 2-MIB synthase, likely representing monoterpenoid cyclases.

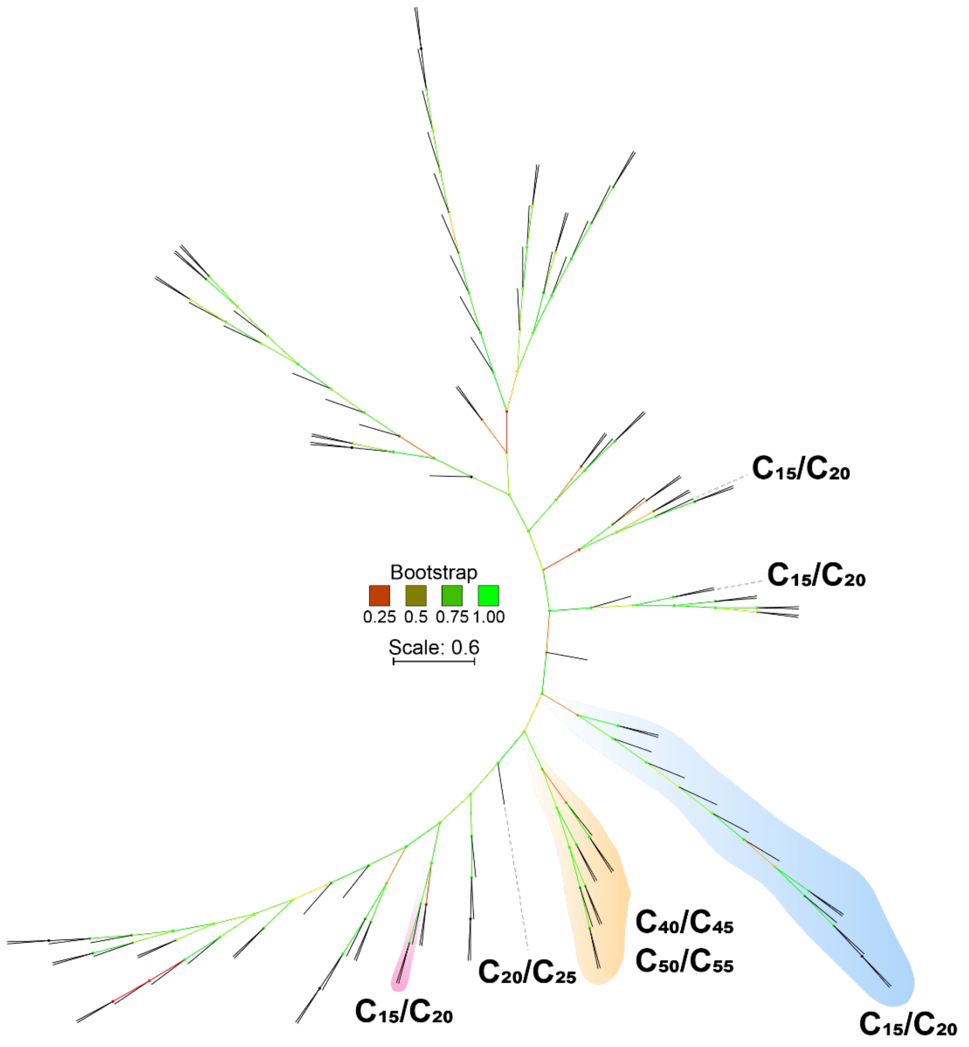

**Fig. S4.** Phylogenetic analysis of 122 Family 2-associated putative polyprenyl transferase (PT) cargo proteins. MAFFT was utilized for multiple sequence alignments with standard parameters; alignment curation was done via BMGE and standard parameters; for tree inference, PhyML+SMS was employed. 25 characterized PTs were included and are highlighted in color. Beginning with dimethylallyl diphosphate (DMAPP), a series of polyprenyl diphosphates are assembled by PTs. Their product range in terms of the length of the synthesized isoprenoid chain is shown in bold. The following nomenclature indicates products with the respective number of carbon atoms in the linear polyprenyl chain: $C_{15}$: farnesyl diphosphate, FPP, $C_{20}$: geranylgeranyl diphosphate, GGPP, $C_{25}$: farnesylgeranyl diphosphate, FGPP, $C_{40/45/50/55}$: polyprenyl diphosphates.

### 3. Additional data and analysis for Family 3 Encapsulins

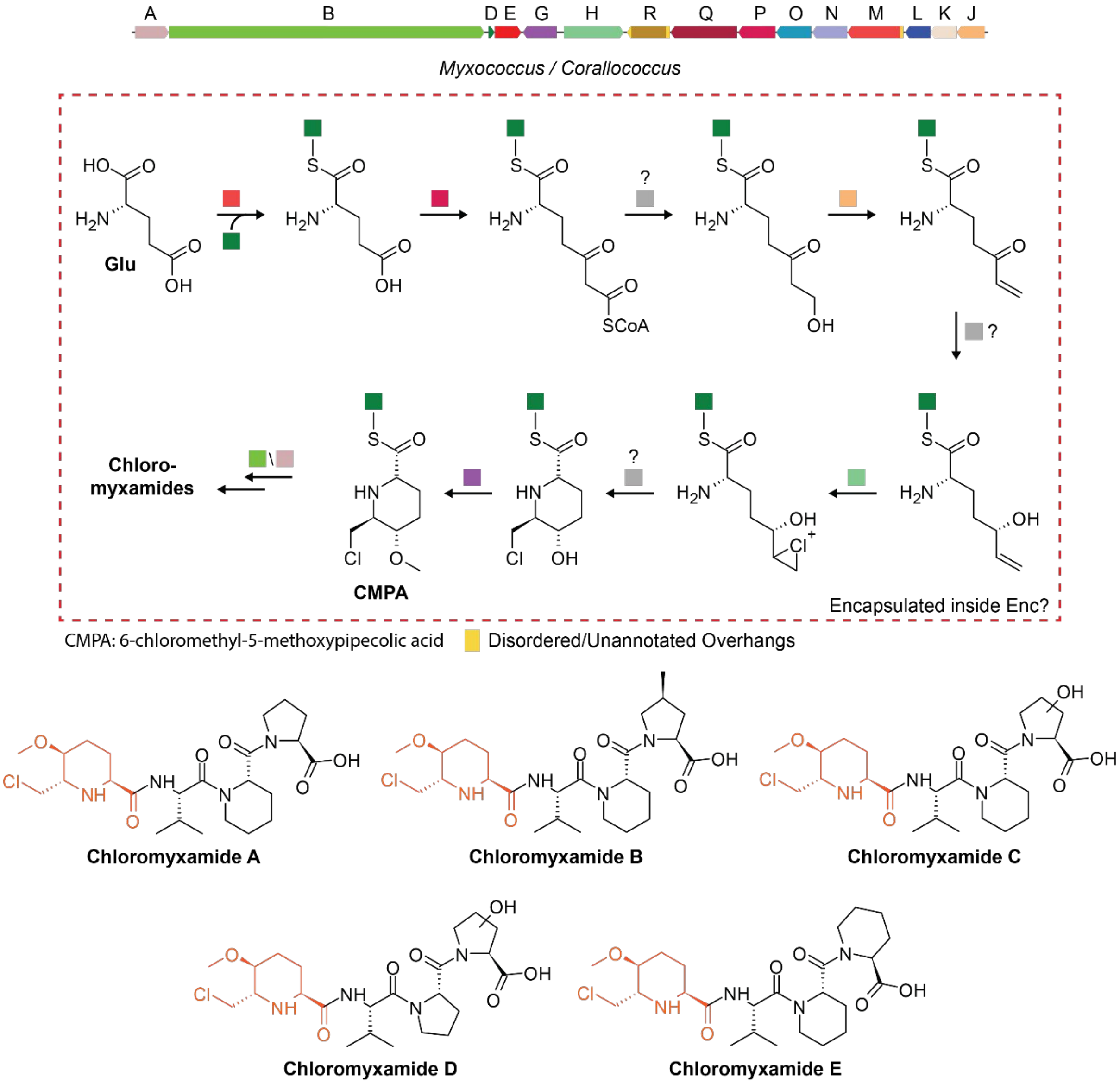

**Fig. S8.** Gene cluster and proposed biosynthesis of chloromyxamides. Top: Family 3 encapsulin-containing chloromyxamide biosynthetic gene cluster and partial proposed biosynthetic route.[4] It is proposed that the CMPA building block is assembled in an amino carrier group-dependent manner. Some or all of the depicted reactions may happen inside a Family 3 encapsulin shell. Disordered and unannotated N-/C-terminal regions of operon components are highlighted yellow and may be involved in mediating cargo loading of select operon components. The chemical logic behind encapsulation may be the sequestration or protection of reactive or toxic aldehyde/ketone or chlorination intermediates. A: ornithine cyclodeaminase, B: hybrid non-ribosomal peptide synthetase/type I polyketide synthase, D: LysW, E: Family 3 encapsulin, G: SAM-dependent methyltransferase, H: rubber oxygenase A, R: aldehyde dehydrogenase, Q: acyl-CoA dehydrogenase, P; acetyl-CoA acetyltransferase, O: acyl-CoA dehydrogenase, N: acyl-CoA dehydrogenase, M: AMP-dependent synthetase, L: NADP-dependent oxidoreductase, K: TetR/AcrR family transcriptional regulator, J: enoyl-CoA hydratase. Bottom: family of chloromyxamides biosynthesized by and isolated from *Myxococcus* sp. MCy10608. The chlorinated CMPA building block is highlighted.

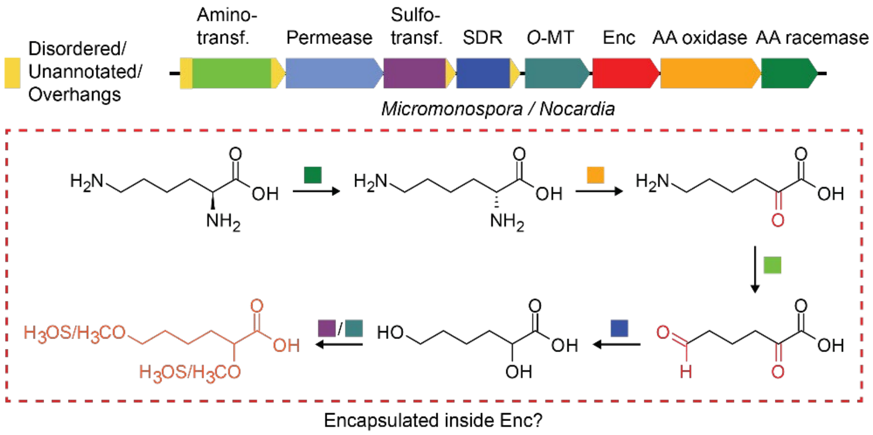

**Fig. S9.** Family 3 operon found in *Micromonospora* and *Nocardia* spp. and proposed biosynthetic route for the encoded natural product. Top: Family 3 encapsulin-containing natural product BGC found in *Micromonospora* and *Nocardia* species. Disordered and unannotated N-/C-terminal regions of operon components are highlighted yellow and may be involved in mediating cargo loading. SDR: short-chain dehydrogenase/reductase, MT: methyltransferase, AA: amino acid. Bottom: Proposed biosynthetic pathway encoded by the operon shown above. Some or all the steps shown may happen inside an assembled encapsulin shell. The goal may be to sequester reactive/toxic aldehyde/ketone intermediates which are highlighted red.

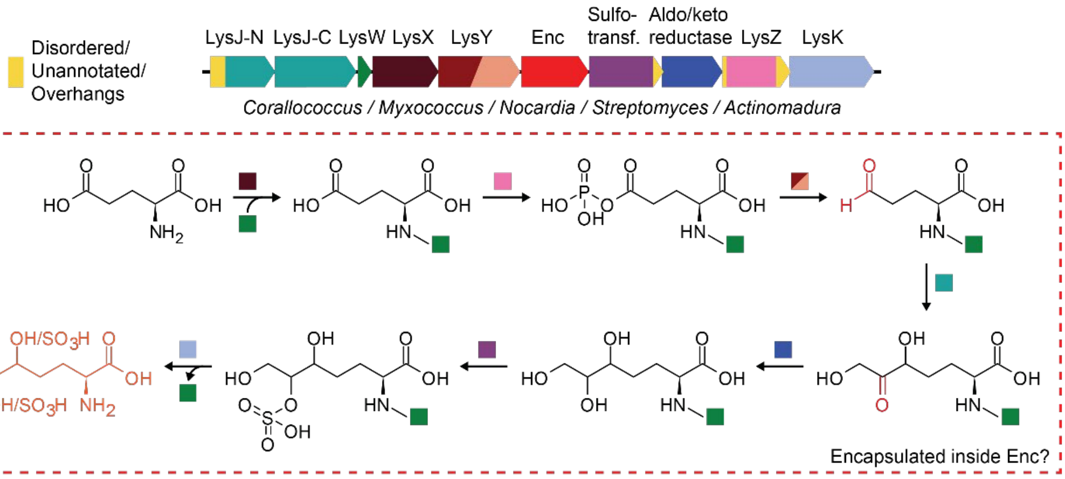

LysJ-N/C: Transketolase LysK: AmCP-AA carboxypeptidase LysW: Amino group carrier protein (AmCP)

LysY: AmCP-AA phosphate reductase LysX: AA-AmCP ligase LysZ: AmCP-AA kinase

**Fig. S10.** Family 3 operon found in many Actinobacteria and *Myxococcus* spp. and proposed biosynthetic route for the encoded natural product. Top: Family 3 encapsulin-containing natural product BGC found in Actinobacteria and Proteobacteria. Disordered and unannotated N-/C-terminal regions of operon components are highlighted yellow and may be involved in mediating cargo loading. Bottom: Proposed biosynthetic pathway encoded by the operon shown above. This pathway seems to rely on an amino group carrier protein to covalently tether pathway intermediates. Some or all the steps shown may happen inside an assembled encapsulin shell. The goal may be to sequester reactive/toxic aldehyde/ketone intermediates which are highlighted red.

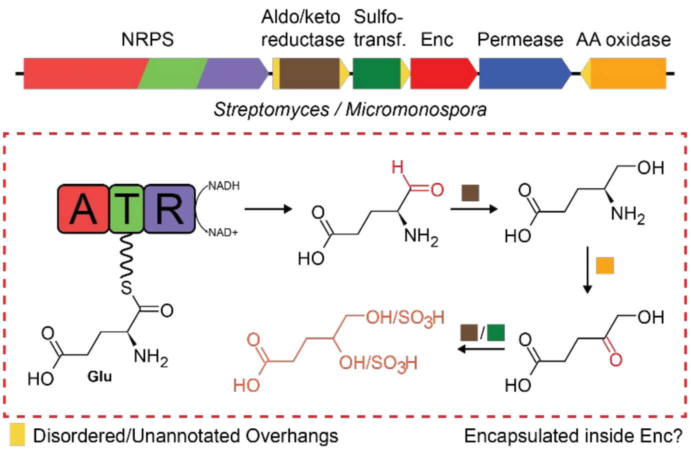

**Fig. S5.** Family 3 operon found in *Streptomyces* and *Micromonospora* spp. and proposed biosynthetic route for the encoded natural product. Top: Family 3 encapsulin embedded in a non-ribosomal peptide synthetase-dependent biosynthetic gene cluster. Disordered and unannotated N-/C-terminal regions of operon components are highlighted yellow and may be involved in mediating cargo loading. Bottom: Proposed biosynthetic route for the gene cluster shown. One or multiple steps may be encapsulated inside a Family 3 encapsulin to potentially sequester reactive or toxic aldehyde/ketone intermediates (red).

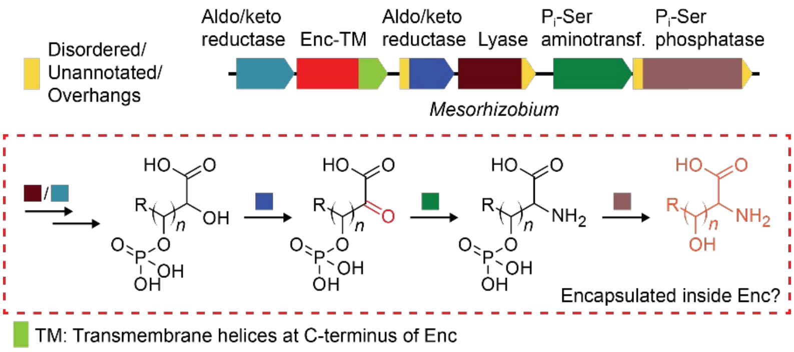

**Fig. S6.** Family 3 operon found in *Mesorhizobium* spp. and proposed biosynthetic route for the encoded natural product. Top: Gene cluster of an unusual Family 3 encapsulin with a C-terminal fusion of hydrophobic/transmembrane helices. Disordered and unannotated N-/C-terminal regions of operon components are highlighted yellow and may be involved in mediating cargo loading. Some of the steps shown might be encapsulated inside this unusual Family 3 encapsulin.

**Family 3 Natural Product encapsulin systems in *Mesorhizobia*:**

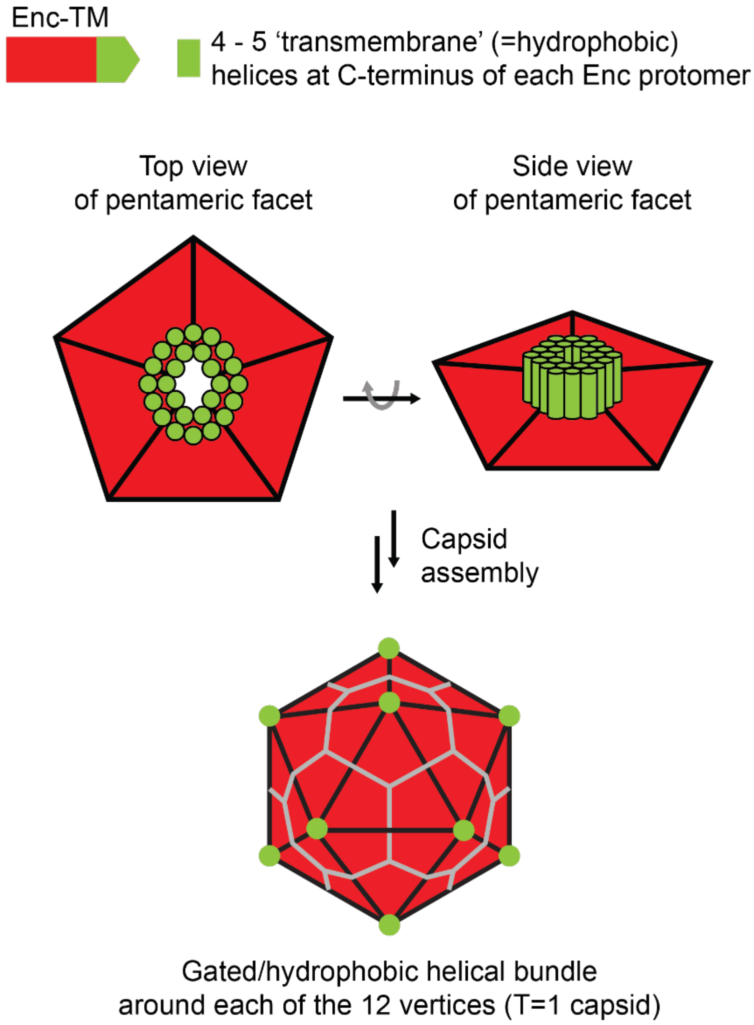

**Fig. S7.** Special type of Family 3 fusion encapsulin found in *Mesorhizobium* species. The conserved HK97-fold encapsulin capsid is C-terminally fused to a 4-5 helix bundle annotated as a major facilitator superfamily (MFS)-type transmembrane protein. Based on the HK97-fold, the C-terminus of the capsid protein is displayed on the outside of the capsid in proximity to the 5-fold symmetry axis/pore. Assuming T=1 icosahedral assembly in the simplest case, this would lead to 5 of these transmembrane or hydrophobic helical bundles meeting at the 12 vertices of the icosahedron. This may lead to the formation of a larger all helical pore-like structure surrounding each 5-fold capsid pore. Assuming this helical bundle behaves in a similar fashion to MFS-type membrane proteins, it is conceivable that these helical bundles sitting atop the pores might act as gates controlling the transport of specific, presumably hydrophobic reactants in or out of the encapsulin shell. This of course assumes that topologically, these helical bundles are inverted compared to MFS transporters, i.e. they are hydrophobic on the inside and hydrophilic on the outside. However, if the helical bundles are hydrophobic on the outside, the same way MFS-type proteins are, it might be conceivable that they are able to directly interact with lipid membranes, potentially anchoring the encapsulin close to or even within a membrane or allow the encapsulin to pass through membranes. A final possibility would be that in analogy to viral envelops, this encapsulin system might be surrounded by a lipid membrane, recruited by the 12 hydrophobic helical bundles displayed at its vertices.

### 4. Additional data and analysis for Family 4 Encapsulins

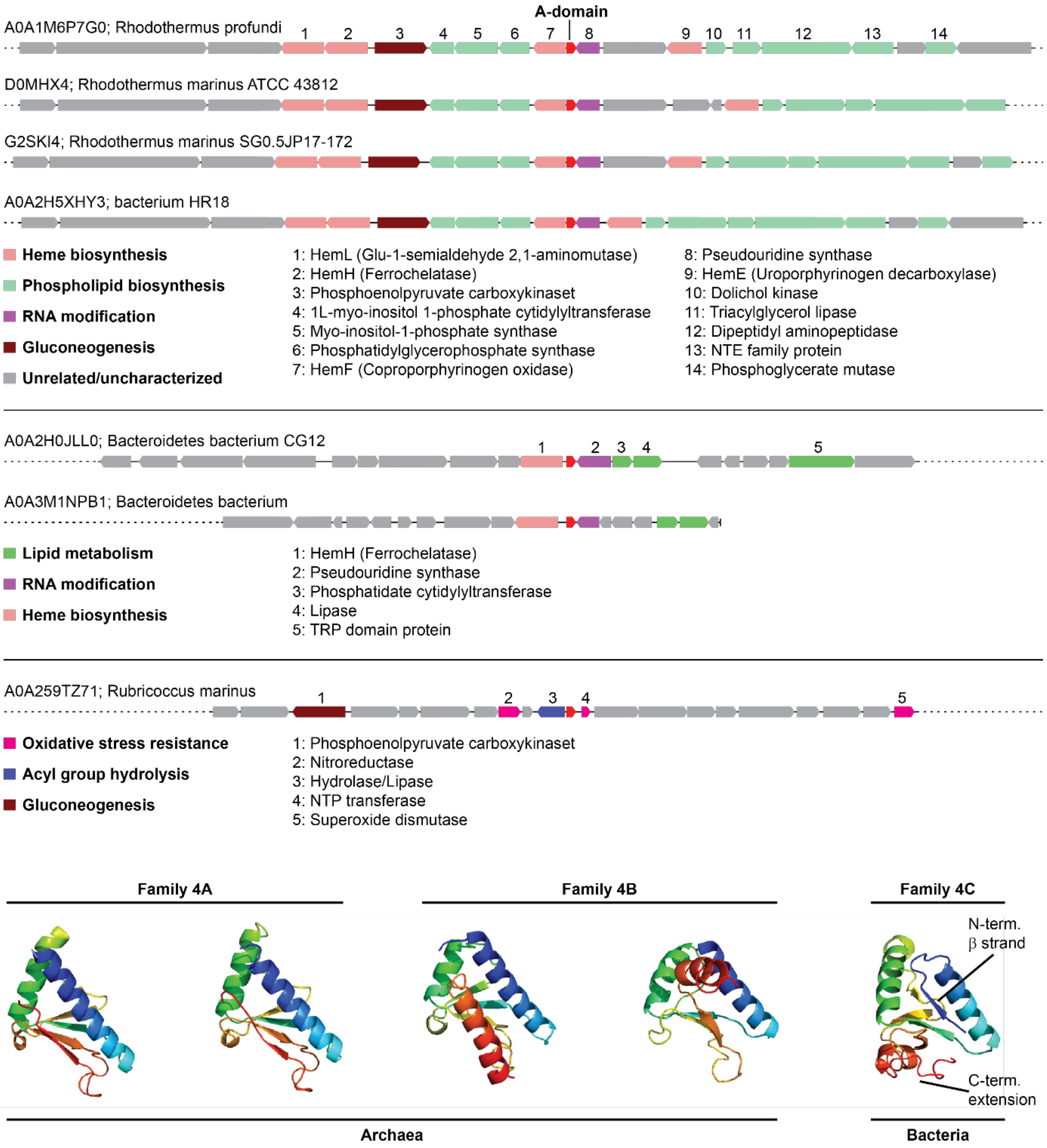

**Fig. S8.** Discovery of bacterial Family 4 systems. Top: Bacterial Family 4 operons. A-domain Encapsulins are not part of an obvious operon structure in the sense of transcription direction making it difficult to propose a putative function. The functional categories of proteins encoded in proximity to bacterial A-domain Encapsulins are highlighted. In particular, heme biosynthesis components seem to be the most abundant components. Bottom: Structural comparison of Family 4A, B and C (bacterial) encapsulins.
